# Division of labor during bacterial warfare

**DOI:** 10.64898/2026.09.21.753294

**Authors:** Divya Choudhary, Youlian Goulev, Carlos Sanchez, Quincey Justman, Johan Paulsson, Kevin R Foster

## Abstract

Division of labor is commonly associated with cooperative behavior, yet one of its most extreme forms occurs during bacterial warfare, where a subset of cells undergoes suicidal lysis to release toxins. Why bacteria divide labor among a few cells rather than producing toxin uniformly remains unknown. Here, we combine timelapse microscopy and simulations, to understand the division of labor during bacterial warfare using bacteriocin (colicin) production by the gut bacterium *Escherichia coli* as a model system. At the single-cell level, we find that lytic toxin production is a tightly regulated event: only cells that commit to lysis produce significant toxin and then lysis only occurs once a large amount of toxin has been made. This high threshold ensures each sacrifice delivers a large dose, which is released in a rapid burst from a lysing cell. While such burst-like release appears to provide no advantage over uniform labor in well-mixed conditions, we show it becomes extremely effective in spatially structured populations where the rapid release of toxin by one cell can generate lethal concentrations locally and eliminate competitors. Finally, we explain why the lysing fraction remains so small. While an increase in the producing cells increases toxin levels, it also increases the probability of local patch extinctions. The division of labor during bacterial warfare, therefore, enables powerful localized killing while safeguarding the population from self-destruction.

## Introduction

The division of labor is fundamental to biological systems where the use of specialized cells or individuals evolves to enhance group functioning^1–5^. Within multicellular organisms, the division of labor promotes cooperation through differentiation into specialized cell types, such as neurons, xylem, and hyphae^6,7^. Even within clonal populations of single-celled bacteria, division of labor creates specialized roles, including motile and sessile cells^8^, enzyme secreting cells^9–11^, biofilm matrix producers^12,13^, or persisters (also a form of bet-hedging)^14^, allowing the population to function as a coordinated collective^3,10,15^. Within societies of organisms, the same logic produces the behavioral and morphological castes of the social insects such as nurse honeybees tending the brood while foragers gather nectar^16^. However, the most striking examples of division of labor arguably arise due to competition and conflict. Famously, honeybee guards suicidally sting mammalian attackers^16,17^. Such a division of labor is also central to the biology of microbes^15,18–20^. Many antibacterial toxins are released through programmed cell lysis of a small fraction of attacker cells ^21–23^, even though bacteria are able to export other toxins without sacrificing the producer^19,24^. Why has this extreme division of labor arisen during bacterial warfare with only a minority of cells lysing while the remainder stay viable? ^4,23,25,26^

The molecular regulators of the production of many antibacterial toxins are well characterised^19,24^, yet the cellular decisions preceding lysis and the impact of toxin released per cell are not. Because lysis is rare and asynchronous, it is challenging to study and has not been followed in single cells over time. Toxin expression is heterogeneous within clonal populations, but it is unknown whether any activated cell can lyse or whether it is a regulated commitment, and therefore self-sacrificial cells differ in amount of toxin released. It is also unclear whether an increase in toxin production in a population occurs via an increase in the toxin released per cell, the number of cells that lyse or both. These dynamics can be particularly consequential for some bacteriocins like colicins in which an initial sacrifice triggers autoinduction of toxin production in neighbors^4,27,28^.

Here we study the production and impacts of colicins^21,29^, plasmid-encoded bacteriocins produced by *E. coli*^28,30^ under the control of the SOS response ^25,31^. We focus on colicin E2, one of the most widespread nuclease weapons in enterobacteria^32^, and common within host gut, where toxin-based competitions can determine which lineages establish^33,34^. Colicin E2 is encoded on a multi-copy colicinogenic (pColE2) plasmid^21,35^, whose operon also carries an immunity protein (ImE2) that neutralizes the nuclease domain of colicin. ImE2 is expressed from both the operon promoter and an independent constitutive promoter^36^, protecting not only colicin-producing cells but also neighboring immune cells not producing colicin from extracellular toxin exposure **[Figure 1A]**. Downstream of ImE2 lies a lysis gene, whose expression causes individual cells to rupture and release toxin into the environment^25,26^ **[Figure 1A]**. Whether such sacrifice events are ultimately beneficial cannot be understood by studying the toxin release alone, because a toxin only matters if it reaches a competitor before it is degraded or lost. In nature, *E. coli* competes in small clumps of cells within dynamic microhabitats (e.g., biofilms^13,19,37,38^, gut crypts^39–43^, and soil aggregates^43–46^), where flow-driven dispersal, crypt flushing, epithelial renewal, or periodic restructuring of aggregates continually remove both the cells and the molecules they release. In such patches, local interactions and diffusion constraints shape competition^13,24,47,48^. Most work on bacteriocins, by contrast, has used millimeter scale colonies or well-mixed liquid cultures in which toxins accumulate over time with minimal loss^21^. These conditions, with many cells present, mask single-cell stochasticity and make the properties and potential benefits of the division of labor difficult to study^48–51^.

**Figure 1:**
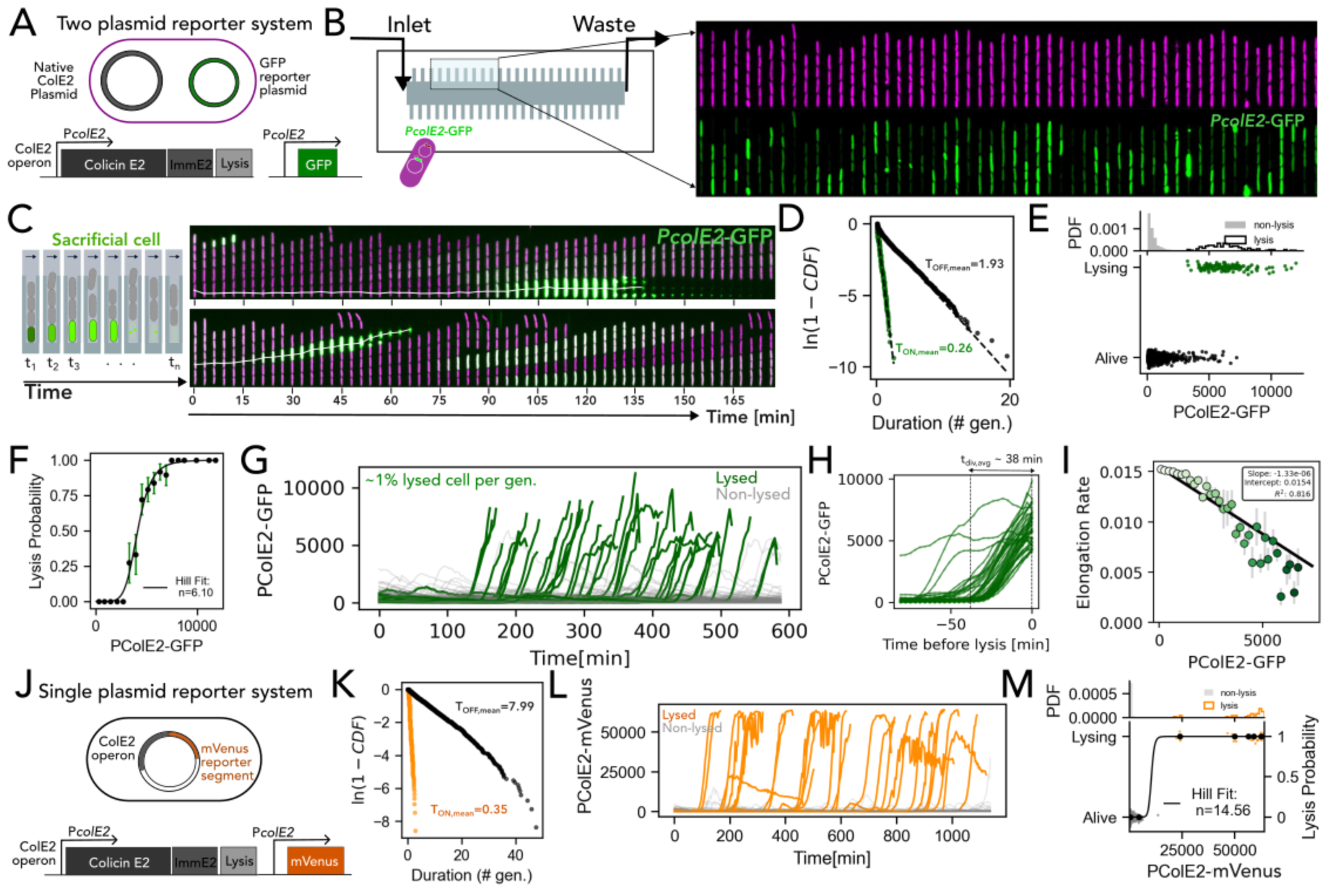
Colicin expression is stochastic, rare, and leads to sacrificial lysis in a subpopulation of killer cells. (A) Schematic of the two-plasmid reporter system with native colicin E2 plasmid and P*colE2*-GFP reporter system. The native pColE2 plasmid encodes the ColE2 toxin, immunity protein (ImmE2), and lysis machinery, while a second plasmid carries a P*colE2*-GFP reporter to monitor toxin production dynamics. (B) Diagram of the ’mother machine’ microfluidic device used for continuous, single-cell lineage tracking, with a representative image showing cells confined to growth chambers. Top shows the constitutive marker (imaged in BFP channel) and the bottom row shows the heterogeneous expression of *PcolE2*-GFP. (C) Representative kymograph of sacrificial cells (marked in white) showing a sharp increase in colicin expression and growth inhibition leading up to lysis (cell death). (D) Log-transformed complementary cumulative distribution ln(1-CDF) of pulse “on” (green) and pulse “off” (black) durations for P*colE2-*GFP. (E) (Top) Probability distribution function of *PcolE2*-GFP expression pulses for alive cells (grey) and cells that lyse (white). (bottom) Expression when *PcolE2*-GFP pulses for alive (black) and lysing (green) cells. (F) Probability of lysis plotted as a function of *PcolE2*-GFP expression pulses for mother cells from panel E. Markers with error bars represent binned experimental data and solid curve shows the corresponding fit to the Hill sigmoid model. The calculated apparent Hill coefficient is displayed in the legend, highlighting a steep switch controlling the commitment to lysis. (G) Fluorescence intensity traces of *PcolE2*-GFP for multiple mother cell (bottom of the channel) lineages over time. A small fraction of cells shows a dramatic spike in fluorescence before lysis (green, ’Lysed cells’), while the majority maintain low expression (grey, ’Non-lysed cells’). (H) Traces of *PcolE2*-GFP for lysing attackers followed from 90 minutes before lysis to assess colicin activation dynamics that precede lysis, dashed line represents average inter-division time. (I) Scatter plot represents median growth rates and *PcolE2*-GFP intensities calculated across discrete non-overlapping bins, with error bars indicating the Standard Error of the Mean. The solid line depicts linear fit to the binned medians, weighted by the inverse of the SEM to prioritize high-confidence data. (J) Schematic of the single-plasmid reporter system, in which an mVenus reporter is placed under the control of the native Pc*olE2* promoter, enabling simultaneous monitoring of colicin expression and cell fate from the native colicin plasmid. (K) Log-transformed complementary cumulative distribution ln(1-CDF) of pulse “on” (orange) and pulse “off” (black) durations for P*colE2*-mVenus. (L) Fluorescence intensity traces of *PcolE2*-mVenus for multiple mother cell (bottom of the channel) lineages over time. A small fraction of cells shows a dramatic spike in fluorescence before lysis (orange, ’Lysed cells’), while the majority maintain low expression (grey, ’Non-lysed cells’). (M) (Top) Probability distribution function of P*colE2-*mVenus pulse expression for alive cells (grey) and cells that lyse (orange). (bottom) Expression when P*colE2-*mVenus pulses for alive and lysing mother cells (orange). The line plot shows the probability of lysis plotted as a function of *PcolE2*-mVenus expression pulses. Markers with error bars represent binned experimental data and solid curve shows the corresponding fit to the Hill sigmoid model. The calculated apparent Hill coefficient is displayed in the legend, highlighting a steep switch controlling the commitment to lysis.

By following individual attackers from activation of the colicin operon to lysis and their neighbors, we track the consequences for the cells around them, using single-cell time-lapse microscopy and microfluidic devices that impose spatial structure and continuous flow. Colicin activation, we find, does not commit a cell to rupture. Lysis is a separate decision, gated at a threshold that promoter strength cannot change, so a lineage can vary how often it lyses a cell but not how much toxin each cell releases. That constraint confines the benefit of dividing labor to the small, rapidly flushed patches in which bacteria commonly compete. We show that a single lysis event is sufficient to eliminate a patch of clonal non-immune neighbors more than ten cell lengths away within minutes. Together, our results establish how rare, programmed cell death enables bacteria to divide labor and mount powerful collective attacks.

## Results

### Colicin E2 release requires surpassing an expression threshold

To investigate the division of labor, we begin by studying the kinetics of toxin gene expression at the single cell level across cells that do and do not commit to toxin release. We resolve cells that commit to toxin release by tracking colicin E2 expression and cell lysis in single cells over many generations in a “mother machine” microfluidic device, which follows thousands of cell lineages over tens of generations^52^ **[Figure 1B]**. We transform a colicin E2 producer (attacker) strain with a low-copy reporter plasmid in which the colicin E2 promoter drives the expression of green fluorescent protein (*PcolE2*-GFP) thereby enabling lineage-resolved colicin expression tracking^4^ **[Figure 1A-B, Movie 1]**. We measure *PcolE2*-GFP expression alongside growth and self-lysis events for each cell at high temporal resolution (generation time = 38 mins) **[Figure 1C, S1A]**.

We observe that colicin activation occurs in pulses with exponentially distributed dwell times in both active and inactive states **[Figure 1D]**, indicating a stochastic process, consistent with previous work over shorter timescales^22,25,27,30^. Further, lysis is rare, with ∼1% cells lysing per generation (*f_lysis_*) **[Figure 1E-G].** What determines which cells lyse? We find that the probability of lysis rises with colicin activation in a nonlinear manner (apparent Hill coefficient, n∼6), above a steep, high threshold (11.1-fold higher than non-lysing cells) **[Figure 1F-G, S2].** Lysis occurs within ∼30 mins of colicin activation **[Figure 1H, Movie 1]**. This sacrificial commitment is restricted to the highest-expressing cells, where rising P*colE2* activity correlates with reduced elongation **[Figure 1I, S1B-C].** These observations support a two-step regulatory model with SOS-driven initiation of colicin production, followed by commitment to lysis above a high threshold of activation, *θ*.

A threshold measured this way is blunted by the measurement itself. In the two-plasmid system, reporter and operon sit on separate plasmids whose copy number fluctuates independently, weakening the correspondence between fluorescence and promoter activation **[Figure 1A]**. As a control, therefore, we built a single-plasmid reporter placing a fast-maturing mVenus^53^ driven by P*colE2* on the ColE2 backbone, so that reporter and toxin experience identical copy-number fluctuations **[Figure 1J, Movie 2]**. Activation remains stochastic with exponentially distributed dwell times **[Figure 1K]**, but the lysis commitment switch is sharper. The probability of lysis rises with an apparent Hill coefficient of ∼15 **[Figure 1M]**, indicating that n∼6 **[Figure 1F]** underestimates the stringency of the threshold. Consistent with this, expression in lysing cells exceeds that of any non-lysing cell by 143.7-fold **[Figure 1L-M, S2].** Because *θ* is far above where the average colicin activation pulses, no cell ruptures until it accumulates a large amount of colicin. This behavior will help to ensure that all lysing cells release a large toxin dose into the environment such that costly cell sacrifice is not spent on low dose releases that are less effective or even non-lethal.

### High toxin expression recruits more sacrifices rather than increasing toxin delivered per sacrifice

The activation threshold *θ* fixes the amount of toxin each lysis releases, and the barrier from baseline expression to *θ* sets how often lysis occurs. At the colicin promoter, that barrier is LexA de-repression ^21,28^. We ask whether *θ* is fixed or scales with overall toxin expression? Doing so addresses whether a lineage raises its toxin expression by sacrifices delivering more toxin or by sacrificing more cells, or both. For this, we construct promoter variants with altered LexA binding affinities (Low, WT, High) in the colicin promoter (P*colE2\**), based on previously characterized mutations^30^. Next, we fuse the promoter variants to colicin E2 (P*colE2*-*colE2) and to mVenus (P*colE2*-*mVenus) on a p15A backbone **[Figure 2A]**. Because LexA represses the colicin E2 promoter, weakening its binding site raises expression and strengthening it lowers expression.

**Figure 2:**
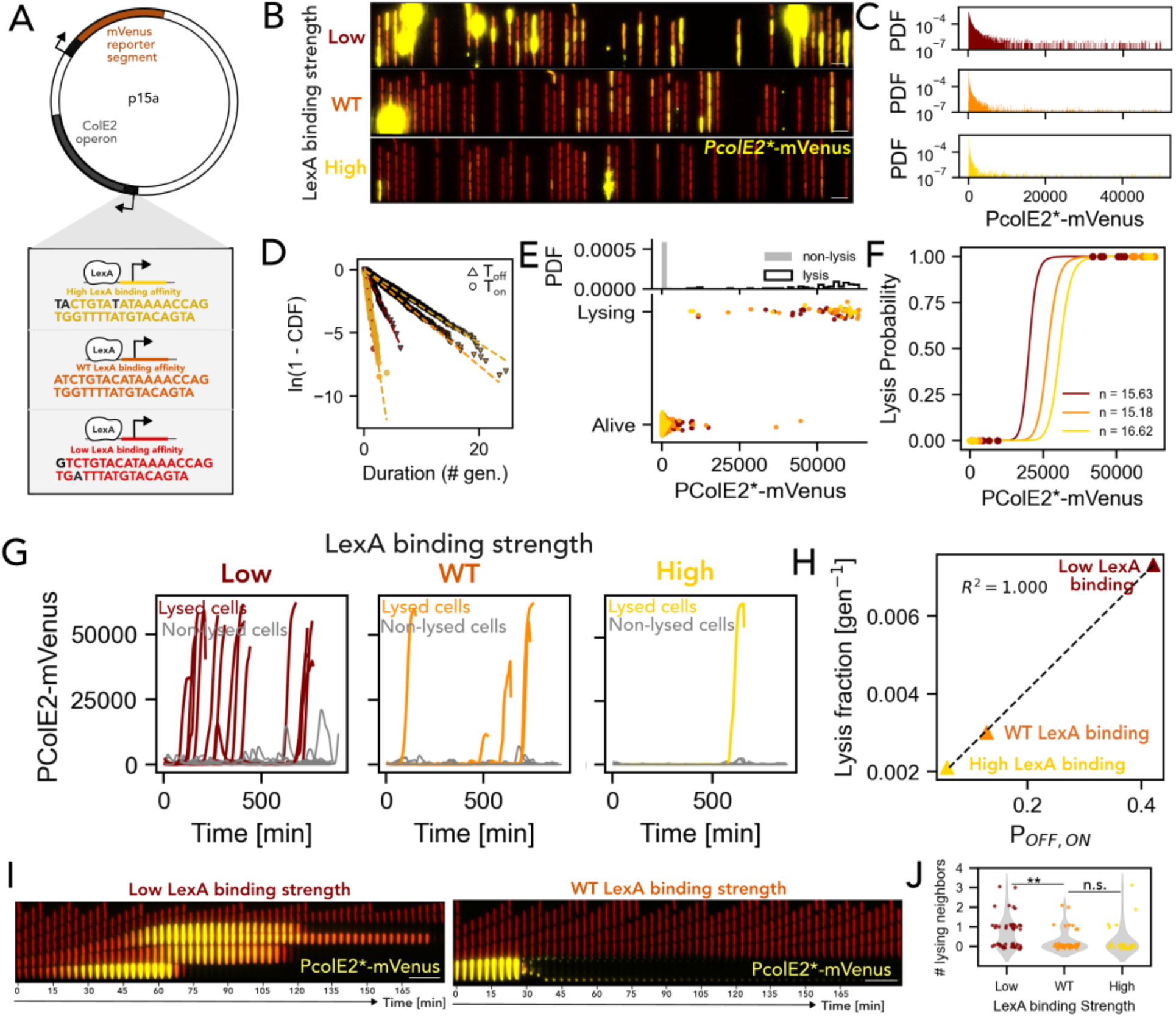
LexA binding strength modulates colicin production and probability for cell lysis. (A) Diagram of the engineered plasmid where the colicin promoter (*PcolE2\**), modified with different LexA repressor binding strengths (High (top), Wild-Type (middle), Low (bottom)), drives expression of colicin and an mVenus fluorescent reporter in distinct transcription units. (B) Representative images of cells with colicin promoter of Low (top), Wild-Type (WT, middle), and High (bottom) strength LexA binding sites, showing that lower binding strength results in higher colicin expression, shown by yellow fluorescence signal. (C) Histograms quantifying *PcolE2\**-mVenus reporter fluorescence for individual cells, confirming the relationship between LexA binding strength and expression levels. (D) Log-transformed complementary cumulative distribution ln(1-CDF) of pulse “on” (circle) and pulse “off” (triangle) durations for P*colE2*-*mVenus for strains with colicin promoter having low (maroon), WT (orange) and high (yellow) LexA binding strength. Best linear fits are shown as dashed lines for each curve. (E) (Top) Probability distribution function of P*colE2*-*mVenus pulse expression for alive cells (grey) and cells that lyse (white). (bottom) Expression when P*colE2*-*mVenus pulses for alive and lysing mother cells for strains with colicin promoter having low (maroon), WT (orange) and high (yellow) LexA binding strength. (F) Probability of lysis plotted as a function of P*colE2*-*mVenus expression pulses for mother cells from panel E. Markers with error bars represent binned experimental data and solid curve shows the corresponding fit to the Hill sigmoid model (strains with colicin promoter having low (maroon), WT (orange) and high (yellow) LexA binding strength). The calculated apparent Hill coefficient is displayed in the legend, highlighting a steep switch controlling the commitment to lysis. (G) Single-cell *PcolE2\**-mVenus fluorescence traces of lysing (colored) and non-lysing (grey) mother cells demonstrate that lower LexA binding strength leads to a higher fraction of sacrificial cells (strains with colicin promoter having low (maroon), WT (orange) and high (yellow) LexA binding strength). (H) The lysis fraction (f_lysis_) plotted against toxin release probability (P_OFF,ON_) for cells with colicin promoter of Low (maroon), Wild-Type (WT) (orange), and High (yellow) strength LexA binding sites. (I) Kymographs following a lysed mother cell for low (left) and WT (right) LexA binding variant. (J) Plot representing number of neighboring attackers that lyse when a mother cell lyses for strains with colicin promoter having low (maroon), WT (orange) and high (yellow) LexA binding strength.

We characterize the LexA variants in the mother machine **[Figure 2B, S3A]**. The low-LexA-binding variant shows a heavier tailed expression distribution and the shortest dwell times in the inactive state **[Figure 2C-D]**. All variants show the same steep dependence of lysis on expression (n ∼ 15) **[Figure 2E-F]**. Critically, the lysis threshold *θ* is similar across the variants, indicating that commitment to lysis requires a fixed amount of colicin, independent of activation rate **[Figure 2E-G, S3B]**. Weakening LexA-binding lowers the barrier to *θ*. For the low-LexA-binding variant, expression in non-lysing cells sits 43-fold below *θ*, compared with 106-fold in WT and 370-fold in the high-LexA-binding variant. A shallower barrier is crossed more often, producing frequent high-expression bursts that surpass *θ* and trigger lysis **[Figure 2G, H]**. Promoter strength therefore sets how many cells lyse but not how much toxin each releases **[Figure 2H]**. A lineage cannot make a sacrifice more lethal, only sacrifice more often, so escalating an attack necessarily means losing more of itself. While other explanations are possible, the invariance in the high threshold *θ* is consistent with a simple model where cells are making and releasing an amount of colicin protein that is close to the physical limits of what a cell can hold in order to maximise the toxicity of each lysis event.

How many cells lyse, however, is not fixed by the promoter sequence alone. Any increase in the SOS signaling lowers the barrier to *θ*, including exposure to colicin released by clonemates, which for colicin E2 mildly induces SOS^27,30^. A lysing cell therefore lowers the barrier in the attackers around it and can pull them across *θ* in turn. We know that *θ* sits so far above the level at which colicin expression typically pulses **[Figure 1L-M, 2E, S2]** that this mild induction is not enough to carry a typical neighbor across it. The number recruited per lysis therefore depends on how many neighbors were near *θ*, which is the quantity the LexA variants manipulate. Weakening LexA-binding raises the fraction of cells sitting close to the threshold **[Figure 2G, S2]**. Tracking neighbors of lysing mother cells, we find coordinated lysis in ∼50% of lysing mother cells in the low-LexA-binding variants **[Figure 2I-J],** but only ∼20% of WT and high-LexA-binding strains. This is consistent with the autoinduction effect in colicin E2 in liquid or agar colony-based competitions^27,30^. Promoter strength thus sets both how often a lineage lyses a cell and how many further cells each lysis takes with it. In the wild type, a lysing attacker recruits fewer than one neighbor, so bursts stay small and self-limiting.

### No benefit of division of labor over uniform labor in well-mixed competition

How efficient are colicin attacks? A minority of colicin producing cells can outcompete colicin sensitive cells in bulk competition^21,23,30^. However, this observation does not by itself explain why colicin release should require a division of labor. For example, it could be even more efficient for all cells to secrete small amounts of colicin, removing the need for sacrificial lysis **[Figure 3A]**. Our single-cell measurements sharpen this problem. Given a lineage cannot make each sacrifice more lethal, only lyse cells more often **[Figure 2]**, the value of the division of labor rests on whether a burst from one cell can achieve a competitive advantage that the same quantity of toxin trickling from many cannot.

**Figure 3:**
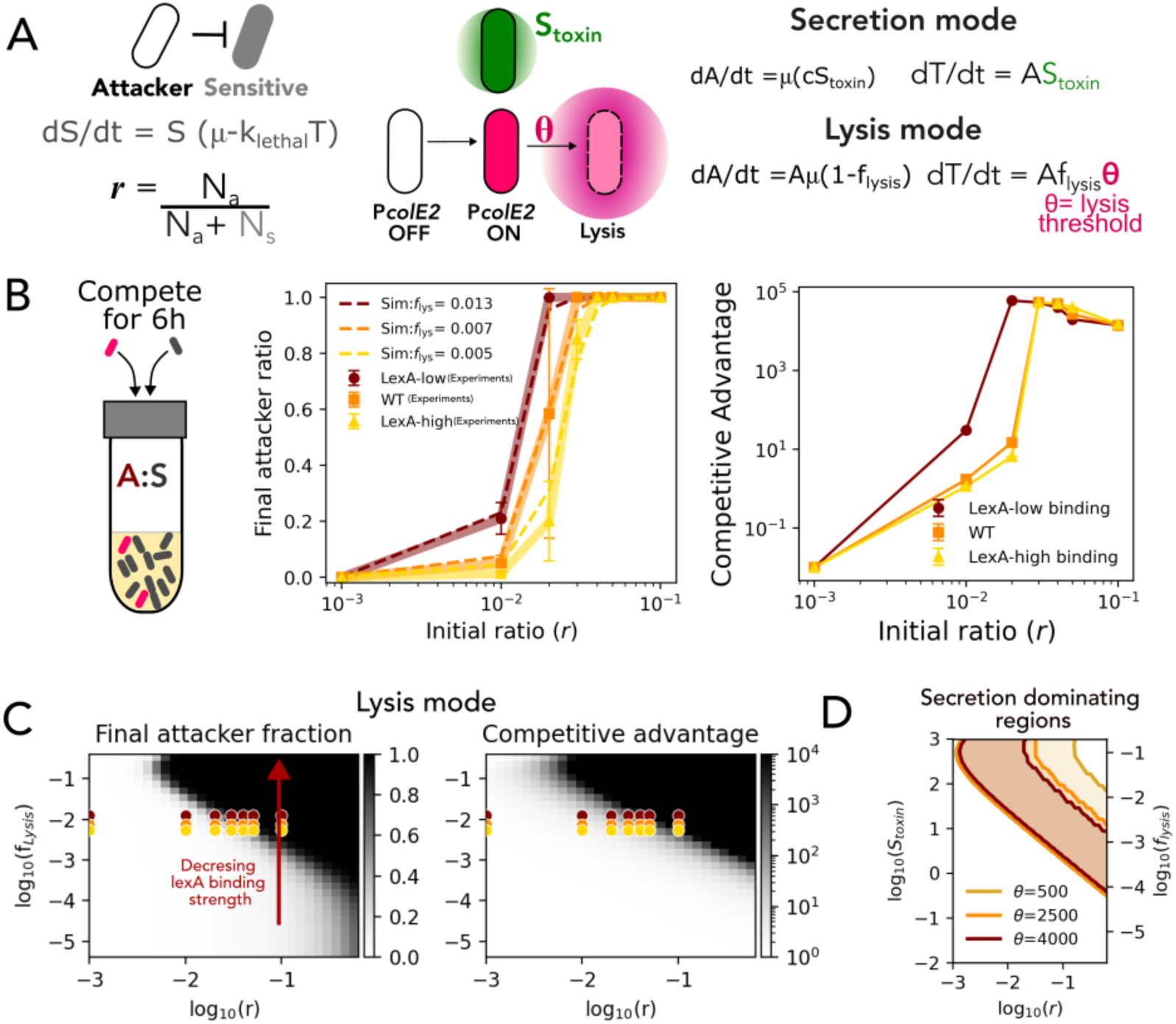
Competitive outcome in well-mixed conditions depends on total toxin, not its distribution. (A) Schematic of the core components for a model simulating competition in well-mixed environments, where an attacker cell either continuously secretes toxin or releases toxin upon passing a lysis threshold in lysis mode. (B) (left) Final attacker ratio plotted against the initial attacker ratio after 6-hour competition between colicin producing and sensitive strain in well-mixed bulk culture. Markers and bold lines show experimentally observed values for mutants with low (maroon), WT (orange) and high (yellow) LexA binding strength on colicin promoter. The dashed line represents the best-fit from simulations generated by optimizing the corresponding fraction of lysis (f_lysis_) values. (right) Competitive advantage of lysing attackers competing in well-mixed liquid culture with susceptible strains, for attackers with colicin promoter driven by LexA of low (maroon), WT (orange) and high (yellow) binding strength across different initial ratios. (C) Heatmap of (left) final attacker fraction and (right) competitive advantage for well mixed competition in lysis mode. The scatter points show the data points represented in Panel B. (D) Contour plots depicting regions where secretion mode is more advantageous (difference in competitive advantage > 0) than lysis mode for different values of lysis threshold (*θ*) when growth penalty is equated. Increasing values of *θ* from yellow to red.

To investigate this question, we develop a minimal model comparing the efficiency of a hypothetical uniform production strategy at rate *S_toxin_* (hereafter called “secretion mode”) with the observed division of labor where lysis of a fraction *f_lysis_* releases *θ* (hereafter called “lysis mode”, **[Figure 1,2]**). To be conservative in looking for a benefit to the division of labor, we assume highly efficient secretion (modeling uniform labor), so that lysis (modeling division of labor) is not simply favored due to limitation in the secretion of proteins. In the model, attackers and susceptible cells are mixed at initial frequency *r* (= number of attackers / total number of cells). Susceptible cells are killed with a rate *k_let_*_ℎ*al*_. In lysis mode, toxin release incurs an explicit fitness cost from sacrificial death, as in the natural case **[Figure 1].** The cost of secretion is modelled as a linear function of *S_toxin_* using the measured growth inhibition relationship **[Figure 1I]**. Here, toxin is distributed instantaneously and evenly throughout the population **[Figure 3A]**. We assess competitive outcomes in two ways: i) final attacker ratio and ii) competitive index, which is based on the number of cell divisions of the competing attacker and sensitive strains **[Figure 3A]**. The latter metric is most representative of fitness under these conditions, but we also include final ratio to allow comparisons with other assays in the paper where this is a better measure of fitness (below).

Because colicin E2 producers release toxin only by lysis, the two modes cannot be compared experimentally, but we can benchmark our modelling predictions by testing how toxin production levels and initial attacker frequency shape competitive outcomes. To do this, we compete a colicin E2 producing strain against an isogenic sensitive strain in shaken liquid cultures (well-mixed conditions) **[Figure 3B**]. We vary the fraction of lysing cells using the LexA variants (P*colE2\**) which vary how often cells lyse (low, a WT and a high LexA binding variant) while keeping toxin released per cell constant, which is the comparison the model requires **[Figure 2]**. Using experimentally measured growth rates together with a fixed killing constant *k_let_*_ℎ*al*_, we estimate the free parameter *θ* by minimizing the error between experiments and modeling predictions. The lysis model reproduces the experimental trends and yields *f_lysis_* consistent with experimentally measured phenotypes **[Figure 3B]**. We find that attacker success depends on *r*, with suppression of susceptible cells occurring when *r* ≥ 3 · 10^−2^. Above this frequency, increasing toxin production does not improve competitive advantage but increases physiological burden by lysing a larger subpopulation. At intermediate *r*, however, weaker LexA binding confers a greater competitive advantage **[Figure 3B**]. In sum, the experiments support the modelling prediction that elevated toxin production only enhances success when attackers are initially rare.

We next simulate competitions across a range of *r* and *f_lysis_*. As expected from the benchmarking, these parameter sweeps show that high *r* and high toxin output (high *f_lysis_*) favor attacker dominance for both strategies **[Figure 3C, S4, S5]**. We next match the costs of toxin production in the two strategies and ask: which mode gives a greater competitive benefit of the attacker cells per unit cost? The toxin released per unit growth cost in lysis relative to secretion is *α* · *θ*, where *α* is the fractional growth penalty per unit toxin. When *α* · *θ* = 1, the two strategies are equivalent. When *α* · *θ* < 1, secretion is favoured because toxin can be produced continuously with less of a cost to population growth rate than cell lysis. Conversely, when *α* · *θ* > 1, lysis wins, that is when the growth cost of secreting *θ* exceeds the loss of an entire cell, which cannot be satisfied in practice. Consistent with this, increasing *θ* reduces the secretion favored region but does not reverse the hierarchy except at 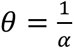, where the two strategies become comparable **[Figure 3D, S4, S5B**]. Raising *θ* makes each death more valuable, but it also makes lysis rarer, and in a well-mixed population the two effects cancel. Thus, in well-mixed conditions, we do not see any scenario in which the division of labor strategy outperforms the uniform secretion.

Outcomes in well-mixed conditions are governed by total toxin production, and not by its distribution across cells. Toxin bursts will be rapidly diluted in well-mixed environments but may be better retained within spatially structured environments that naturally arise in many natural systems^13,19,54,55^. For example, in the mammalian gut where *E. coli* commonly resides^39–42,56^, peristaltic mixing is intermittent and there is considerable spatial structure due to food particles in host anatomy, including blind ended crypts where cells exist in dynamic small clumps and toxins have the potential to act locally. We hypothesized, therefore, that this spatial structure has the potential to influence the value of releasing toxins from only a few cells as compared to the majority of cells.

### Competition for a dynamic local patch favors the division of labor

To capture competition in spatially structured environments, we simulate cells and toxin spread explicitly in space and time **[Figure 4A]**. Cells occupy discrete sites on a two-dimensional lattice and grow within a closed-end chamber **[Figure 4B]**. Toxin released by a focal attacker diffuses away rather than mixing instantly **[Figure 4A-B]**. We again compare the two modes of toxin production, i) uniform secretion, and ii) lysis at equal net growth cost. We assess competitive success by strain frequency, which captures the ability of a strain to become numerically dominant and take over the patch. Under such intense local competition for space, this metric is likely to be more representative of ecological and evolutionary success than a metric based on cell divisions **[Figure 3]**. This metric is also most relevant because the highest fitness strategies are expected to be those that can take over a patch and then shed dispersing cells over a long period, which may not be the same strategies that display the fastest growth rate, particularly early on in the model.

**Figure 4:**
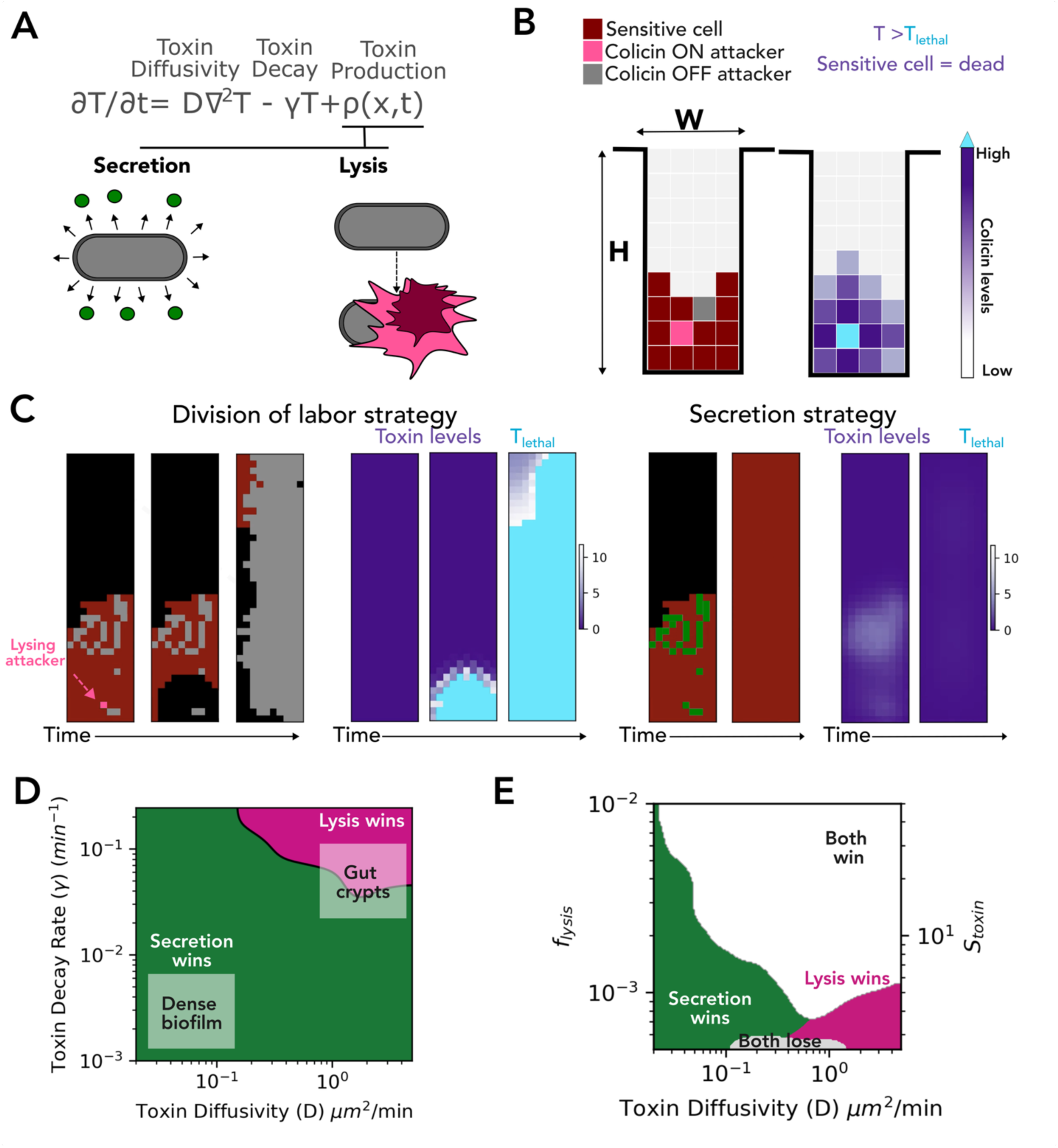
Lysis strategy dominates in structured environments with flow. (A) Incorporating the diffusion of toxin in spatial scale. (B) Schematic of the 2D lattice model where toxin diffuses locally from a lysing cell. (C) Representative snapshot of modeling results for (left) division of labor strategy and (right) secretion strategy starting with the same initial condition. (left) Grey points depict attackers (green if lysing in the next frame) and maroon points depict sensitive cells over time while competing. (right) Toxin levels with higher levels depicted in lighter shade of purple and toxin levels > *T_let_*_ℎ*al*_ are shown in cyan. (D) Heatmap of difference in final attacker fraction for lysis strategy (division of labor, purple) versus secretion (uniform labor, green) strategy for spatial competition with different values of toxin diffusivity (D) and toxin decay rate (*γ*) starting with initial attacker ratio of 0.1. At low D and low *γ*, representing niches like dense biofilm, secretion dominates. However, for high D with high *γ*, representing dynamic niches like gut crypts or soil aggregates with high turnover in space, lysis strategy dominates. (E) Heatmap of difference in final attacker fraction for lysis strategy (pink, division of labor) versus secretion (green, uniform labor) strategy for spatial competition with different values of D and toxin production values (f_lysis_ and corresponding S_toxin_). At high toxin production both strategies dominate. At low D and high toxin production, secretion dominates. However, for high D with low toxin production, lysis strategy dominates.

Across toxin diffusivity (D) and toxin decay (*γ*), the strategies diverged in competitive advantage **[Figure 4C-D, S6]**. At low D and low *γ*, representing conditions of a dense matrix, secretion won by sustaining a stable lethal zone where a lysis burst could not diffuse far enough to kill competitors. At high D and high *γ*, such as in high-turnover environments subject to advective flow or high protease activity (e.g. gut crypts), diffusion and decay outpace slow toxin accumulation, rendering steady-state secreted toxin levels ineffective for killing. In these high-clearance regimes, sacrificial lysis becomes an effective competitive strategy **[Figure 4D]**.

Diffusion and decay are set by the habitat. What a lineage controls is the toxin output, so secretors could simply compensate by producing more. We, therefore, compare both strategies across a range of *S_toxin_* or *f_lysis_* and toxin diffusivity D. At high toxin production *S_toxin_* or *f_lysis_* (or high *r*), both strategies are effective [**Figure 4E, S6]**, but the strategies diverge at low toxin output. There, secretion fails to reach local lethal threshold at high diffusivity [**Figure 4E]**, whereas stochastic lysis of a minority functions as a pulsed release, transiently spiking local toxin concentrations above killing threshold and eliminating adjacent competitors. The surviving attacker cells then expand into cleared sites. Notably, this competitive advantage occurs where the well-mixed model predicted **[Figure 3C]**, and experiments confirmed **[Figure 3B]**, that attackers could not eliminate susceptible cells. In bulk, a lysing cell contributes its toxin to a population large enough to dilute it but in a patch of tens of cells, the same toxin saturates the neighborhood. These results lead us to a key testable prediction: colicin-producing cells should become effective competitors in structured environments with continuous flow, even when rare.

### Spatial confinement reverses the well-mixed prediction and enables rare attackers to dominate

To impose spatial structure with continuous environmental turnover, we develop a microfluidic device, the “family machine”, designed to mimic structures like blind-ended gut crypts^39–42^. A main flow channel connects orthogonally to an array of wider growth chambers (dimensions: 25 *μ*m X 100 *μ*m), allowing cells to expand in two dimensions while remaining exposed to advective flow **[Figure 5A-B]**. Each chamber holds a small, self-contained population (∼100-200 cells) under constant clearance, reproducing the two features our modelling identifies, which are spatial structure and turnover fast enough that toxin cannot accumulate. We compete a colicin E2 producer (unlabeled, attacker), against a sensitive strain constitutively expressing *PrpsL*-mKate2 and carrying a (*PrecA*-sCFP3A) reporter, allowing DNA damage by colicin attack to be visualized in real time **[Figure 5B-D, S7, Movie 3]**.

**Figure 5:**
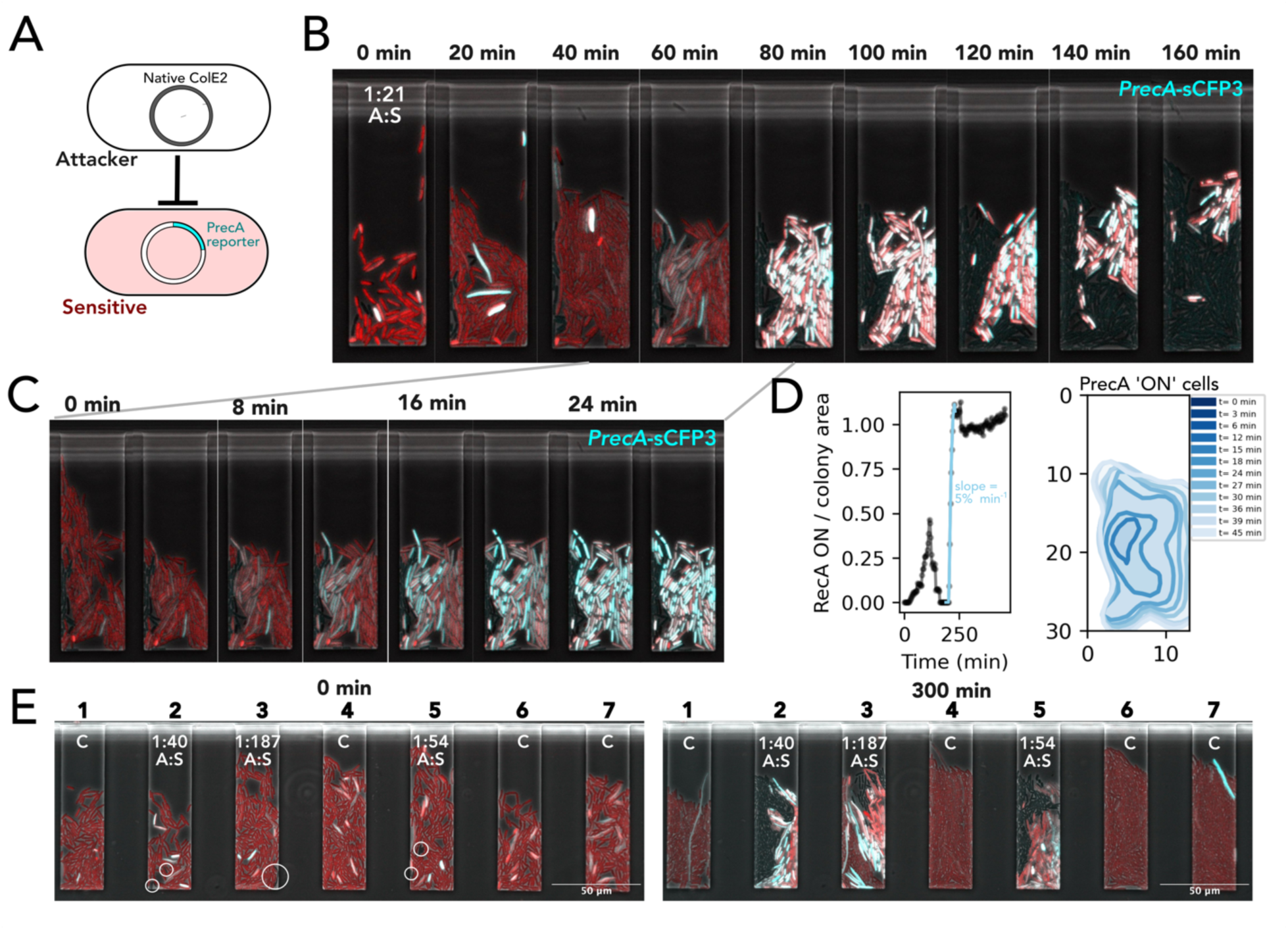
Colicin-mediated killing in spatially structured patches under diffusion constraints. (A) Schematic of the competition assay in microfluidic growth chambers. Unlabeled colicin-producing ’Attacker’ cells are co-cultured with ’Sensitive’ cells expressing a fluorescent reporter for DNA damage (*PrecA*-sCFP3A). (B) Kymograph of a single growth chamber. Sensitive cells (red) are progressively killed following the induction of the DNA damage reporter (cyan), indicating colicin activity. (C) A high-temporal-resolution view of the onset of killing, showing the rapid spread of DNA damage signaling spatially among sensitive cells. (D) (left) Fraction of chamber colonized by sensitive cells that is P*recA* ‘ON’ over time for traces in panel B, the blue points show the time when colicin released by lysing attackers induces DNA damage in sensitive cells. (right) The spatio-temporal spread of P*recA* ‘ON’ cells during the time depicted in blue in the left plot. (E) Snapshots of seven parallel microfluidic chambers at the start of the experiment (0 min) and after 300 minutes. While chambers are initially populated with a mix of attacker (circled, black) and sensitive cells (red, P*recA* induction in blue), they are later dominated by the surviving lineage, demonstrating patch takeover. Lineages with even a few attacker cells are dominated by growing attackers and growth arrested, damaged sensitive cells.

Chambers containing only sensitive cells showed no RecA induction, confirming that killing resulted from local colicin activity within the chamber and that toxin did not transfer between chambers **[Figure 5E]**. In mixed chambers with both attacker and susceptible strains, in contrast to well-mixed conditions **[Figure 3]**, attackers dominate even when initially rare (*r* ≪ 3 · 10^−2^). We find good agreement, therefore, with our spatial modeling predictions **[Figure 4, 5E]**. DNA damage followed by P*recA* fluorescence and growth arrest originate adjacent to the attacker cells and propagate over micrometer distances within minutes **[Figure 5C-D, S7]**. In this way, toxin from only a few sacrificial attackers reaches an entire chamber-sized patch and triggers a spatially spreading wave of intoxication in sensitive neighbors **[Figure 5E]**. The attackers then proliferate to occupy the patch demonstrating the remarkable ability of these bacterial toxins, even when released from a few cells, to be decisive for ecological outcomes. A lineage carrying a rare, stringently gated suicide program can therefore take over a patch from a frequency at which bulk competition predicts its elimination.

### Increased recruitment into lysis risks lineage collapse

If toxin released from a sacrificial lysis is so effective, why do bacteria allocate so few cells to it? Intuitively, one might expect that there would be an even greater benefit if more cells were allocated to produce toxin. To address this, we return to the *PcolE2*-GFP reporter characterized in the mother machine **[Figure 1]**. By co-culturing *PcolE2*-GFP labeled attackers with the sensitive strain, we spatio-temporally follow toxin activation, lysis, target intoxication and killing in a single experiment. We find that induction of *PcolE2*-GFP in just a single attacker is sufficient for mass killing of sensitive cells within minutes of lysis **[Figure 6A-D, Movie 4]**. A single death therefore already allows attackers to dominate, and additional sacrifices in the same patch add little to the competition. What they do add is the risk of having too many cells lyse in a patch and undermine the ability to colonize. This risk is particularly high when bacteria use a coordinated attack as occurs with bacteriocins like colicin E2, where release in one cell can trigger lysis in neighboring cells via autoinduction. Our data shows that colicin production increases in attackers neighboring a lysing attacker for ∼ 30-40 minutes, while the sensitive cells are being eliminated **[Figure 6A, C]**, allowing more cells to surpass the lysis threshold, generating a coordinated burst. **[Figure 6A]**. Recruitment is the only route by which a lineage can concentrate more toxin into a single moment, since *θ* fixes the amount of colicin that each individual lysis delivers **[Figure 2E-G]**.

**Figure 6:**
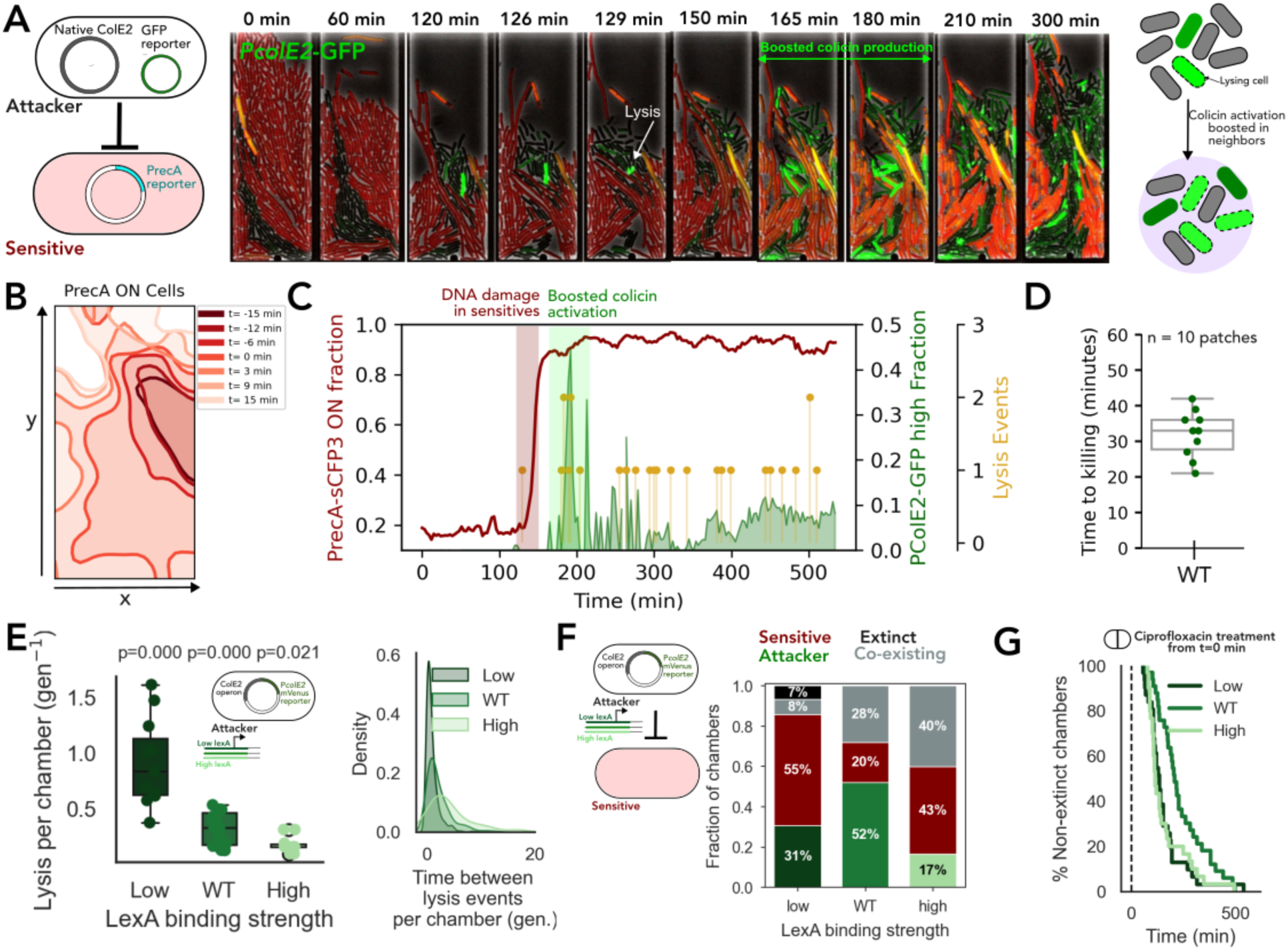
A single attacker can recruit neighboring attackers amplifying attack. (A) (left) Schematic of the colicin expression reporter strain (*PcolE2*-GFP) in unlabeled attacker cell. (middle) Kymograph shows colicin producing (green) attacker cell (marked with arrow) lysed at 129 minutes. The attackers expand into the space cleared of sensitive cells (red). (right) Schematic representing the autoinduction effect at the individual cell level in patches, where local release of toxin by a lysing focal cell boosts toxin production in nearby attackers leading to lysis of those that had a higher colicin activation level at the time of focal cell lysis. (B) The spatio-temporal spread of RecA ‘ON’ sensitive cells from 15 minutes before and after lysis depicted in panel A. (C) Kinetics of a localized colicin attack in Panel A tracked over 500 minutes in a microfluidic chamber. (left axis) Fraction of area occupied by sensitive cells exhibiting active SOS response, tracked with the P*recA* transcriptional reporter. The red shaded window highlights the abrupt onset of widespread DNA damage in sensitive cells following one lysis event releasing colicin. (Right axis, green) Fraction of area occupied by attacker cells activating a high P*colE2*-GFP expression. The green shaded window highlights a wave of boosted colicin activation recruited locally by the first lysis event. (Right, yellow) Discrete, single cell autolysis events plotted over time, The clustering of yellow dots inside the green activation window demonstrates the transition to coordinated colicin attack. (D) Boxplot represents the time between detection of attacker cell lysis and the death of sensitive cells in 10 individual chambers. (E) Lysis dynamics for strains with different LexA binding strengths in microfluidic growth chambers. (left) Number of lysis events per chamber per generation. (right) Distribution of time interval between subsequent lysis events within a chamber. (Schematic shows variants with colicin promoter with low (darkgreen), WT (green) and high (lightgreen) LexA binding strength). (F) Quantification of competition outcomes from experiments for ∼250 growth chambers per strain and 3 repeats. Only chambers initially containing both strains were analyzed, and outcomes were classified as sensitive strain takeover (maroon), attacker strain takeover (green), both strains co-existing (grey), both strains extinct (black). The WT strain (intermediate production) shows the highest frequency of successful patch takeover by attackers. (G) Percentage of non-extinct chambers when colicin producing variants were exposed to ciprofloxacin treatment at t=0 minutes. Variants with colicin promoter with low (darkgreen), WT (green) and high (lightgreen) LexA binding strength

Coordinated toxin release can, therefore, be effective for rapidly eliminating a competitor. However, why do strains not lyse a larger fraction of cells to release even more toxin? To address this, we compete LexA variants **[Figure 2]** against sensitive cells. In attacker-only chambers, the low-LexA-binding variant lyses most often (0.914 +/- 0.119 events per generation per chamber), roughly three-fold more than WT (0.310 +/- 0.035) and five-fold more than the high-LexA-binding variant (0.176 +/- 0.024), with correspondingly shorter intervals between successive events (1.15 +/- 0.09 generations, versus 3.36 +/- 0.29 and 6.87 +/- 1.11) **[Figure 6E, S8, Movie 5]**. Clustered intervals are the signature of recruitment rather than of independently activating cells.

In competition against sensitive strains **[Figure 6F]**, the WT clears its competitors and colonizes 50% of the chambers despite its intermediate lysis frequency, consistent with a single lysing cell clearing the surrounding patch of sensitive cells **[Figure 6F]**. The high-LexA-binding strain lyses too rarely to produce sufficient colicin and colonizes 17% of the patches, indicating too extreme of a division of labor. More surprisingly, the low-LexA-binding strain also performs poorly, colonizing 31% chambers. In addition to the cost of losing cells to lysis and slower population growth, 7% of competitions caused the low-LexA-binding strain to drive both itself and the susceptible strain to extinction within a chamber **[Figure 6F, S9]**. Because a cell sits far below *θ*, the mild SOS induction that follows a neighbor’s lysis allows only the few cells close to the threshold to lyse. A population with more cells near *θ* as in low-LexA-binding strain would sweep many neighbors into lysis and cost the lineage the patch it had just cleared.

Finally, we asked whether external DNA damage imposes the same constraint. Ciprofloxacin is a DNA-targeting antibiotic that induces the SOS response, thereby de-repressing the colicin promoter. Local colicin E2 release from lysed cells can therefore prime neighboring cells for incoming damage, but excessive SOS activation can instead trigger runaway lysis and chamber extinction. Following ciprofloxacin exposure, WT populations went extinct more gradually than either the low- or high-LexA-binding variants, consistent with an intermediate level of priming that enhances the response without triggering widespread lysis **[Figure 6G, Movie 6]**. In contrast, the low-LexA-binding strain showed coordinated lysis, such that the uniform SOS signal drove a large fraction of cells into lysis within minutes. Thus, while lysis can provide a competitive advantage, excessive synchronization of lysis can eliminate the population itself. Stochastic local extinction would also become a risk for non-autoinducing strains above some lysis frequency, such that extinction risk is likely to be a general constraint whenever bacteria compete in small patches with low cell numbers.

## Discussion

Many species of bacteria use division of labor to release toxins that kill or inhibit competing strains, where only a few cells release the toxins^18,19,21,30^. Here we set out to explain why bacteria have evolved such an extreme division of labor during competition. Colicin E2 activation in individual cells is stochastic, but we find that the lysis is not its automatic consequence. Commitment to lysis requires crossing a sharp threshold in colicin expression, such that only a rare subpopulation ruptures. Because this threshold sits far above the average colicin expression, a lysing cell carries more than 100-fold more toxin than any survivor, and the release is concentrated into brief, large bursts. The threshold is a fixed property of the operon, unchanged by promoter strength, so a lineage cannot make each sacrifice more lethal by modifying gene expression. It can only make sacrifices more often. Where this strategy pays off depends on the environment. In well mixed conditions, we found no clear advantage of the lysis strategy over a simpler mode in which all cells continuously secrete small amounts of toxin, because outcomes there depend more on the total toxin produced than on how it is delivered. Under spatial confinement, however, this burst release becomes advantageous because toxin accumulates locally before it is diluted or removed, so a single lysing cell clears neighboring competitors within minutes and opens space for expansion. This allows attackers to take over a patch from frequencies at which bulk competition predicts their elimination. The advantage of this strategy is strongest in environments where toxin decays rapidly, for example due to advection and fluid flow in gut crypts or other flow-exposed habitats, conditions under which continuous secretion can never sustain a lethal zone.

Because the lysis threshold is fixed, a lineage produces more toxin by sacrificing more cells. Toxin released by one cell enters the neighboring clonemates and lowers their barrier to cross the threshold, so increasing the frequency of lysis raises both how many cells reach the threshold and how many further cells are pushed across the threshold to lyse with it. Our LexA variants further show that lysis frequency must be finely tuned. Weakening LexA repression triples lysis and increases the fraction of lysis events accompanied by a neighbor’s lysis. While low-LexA-binding variants enhance attack, excessive lysis becomes self-defeating and collapses the attacker population itself. Strong repression, in contrast, suppresses toxin release too much. The wild-type promoter, therefore, appears to occupy an intermediate regime that effectively balances killing with preservation of survivors.

How general are our findings? Although we focus on the specific case of *E. coli* colicins, the principles identified here are likely to apply more widely. Many bacterial species release bacteriocins by lysis^21,57,58^, including homologues of the colicins^29^ in *Pseudomonas aeruginosa*^59^, *Salmonella enterica*^60,61^ and *Klebsiella pneumoniae*^62^. Cell lysis is also used to release bacteriocins that are unrelated to colicins, including phage-tail-like bacteriocins (tailocins)^63,64^ and the TcdE-dependent lysis cassette of *C. difficile*^65^, which are derived from phage. This strategy also extends to major human pathogens, for example, increased pathogenicity of *Streptococcus pneumoniae* (serotype 1) is driven by rapid autolysis, which releases pneumolysin into the host environment^66,67^. In cases where release is destructive and the toxin released per cell is fixed, the same trade-off between the value of cell death and its frequency should apply.

The benefit we observe for division of labor during bacterial warfare rests on spatial structure and effective diffusion, and our modeling predicts where it should be effective. This includes environments in which toxin is cleared fast enough that continuous secretion cannot sustain a lethal zone yet retained long enough for a burst to act. Strongly diffusion-limited environments fall outside this window, where a dense matrix prevents the rapid dispersal of toxin from lysed cells^13,19,37,38^, continuous secretion is the better strategy. Constant mixing and renewal in the upper layer of the gut crypt allows for toxic molecules to persist locally long enough for lysis events to concentrate activity near the epithelium^39,40,56,68–71^. Similar conditions are likely to occur elsewhere in the mammalian gut where food particles and other host features generate structure and in soils, where antimicrobial-producing bacteria reside^46^ and metabolites diffuse through thin water films surrounding soil aggregates^45,55,72^.

Our experiments also underline that toxin release is highly plastic^30,73–75^. The colicin E2 operon is governed by the SOS response^28,30^, and cell-cell variation in spontaneous DNA damage generates the extreme division of labor. In addition, this regulatory wiring acts as a mechanism of competition sensing^76^. Exposure to exogenous DNA damaging bacteriocins can trigger a coordinated counterattack, including mass cell suicide in biofilms where toxin accumulates strongly because decay is minimal^4,27,30^. In the inflamed gut, host-derived DNA-damaging agents and high-density blooms can further amplify autoinduction ^60,77,78^. More generally, by coupling toxin release to global stress networks, bacteria can dynamically tune their division of labor, scaling from rare local sacrifice to population-wide warfare according to the severity of the ecological threat^76^.

Division of labor in bacteria is well known as a way to facilitate cooperation among cells^3,8,10–12,15,19,79,80^. Our results show that the division of labor is also extremely effective for sets of cooperative bacteria to perform aggressive functions. When bacteria compete locally, concentrating toxin release in just a few cells is decisive for ecological outcomes.

## Acknowledgments

We thank Somenath Bakshi, Maxence Vincent, and members of the Paulsson lab for their discussions and comments on the manuscript. Research in the Paulsson lab is funded by ARPA-H. D.C is supported by an HFSP postdoctoral fellowship. We thank the Foster lab for sharing the BZB1011 strains harboring P*colE2*-GFP and ColE2 plasmids.

## Author contributions

Conception and design of study, D.C., and K.R.F., engineering of genetic constructs, D.C.; microfluidic setup, D.C., and C.S., mother machine chip design and fabrication, D.C., and Y.G., data analysis and interpretation, D.C., J.P., and K.R.F.; writing and editing of the article, D.C., Q.J., J.P., and K.R.F.

## SUPPLEMENTARY INFORMATION

### Supplementary figures

### Supplementary Movies

***Movie 1.*** Time lapse of P*colE2* promoter expression (P*colE2*-GFP, green) overlaid with constitutive expression of BFP (magenta) for BZB1011 growing in microfluidic trenches. The reporter is carried on a separate plasmid from the native pColE2 plasmid. Scale bar = 10 μm.

***Movie 2.*** Time lapse of P*colE2* promoter expression (P*colE2*-mVenus, yellow) overlaid with constitutive expression of P*rpsL*-mKate2 (red) for MG1655 growing in microfluidic trenches. Here the reporter is carried on the pColE2 plasmid itself, so reporter and toxin experience the same copy-number fluctuations. Scale bar = 10 μm.

***Movie 3.*** Time lapse of competition assay in three microfluidic growth chambers. Unlabeled BZB1011 colicin-producing attacker cells are co-cultured with colicin-sensitive cells (maroon) expressing a fluorescent reporter for DNA damage (*PrecA*-sCFP3A, cyan).

***Movie 4.*** Time lapse of competition assay in ten microfluidic growth chambers. Unlabeled BZB1011 colicin-producing attacker cells expressing a fluorescent reporter for colicin expression (P*colE2*-GFP, green) are co-cultured with colicin-sensitive cells (maroon) expressing a fluorescent reporter for DNA damage (*PrecA*-sCFP3A).

***Movie 5.*** Time lapse of colicin E2 producing MG1655 strains with colicin attackers (red) with (left) Low, (middle) WT, and (right) High LexA binding strengths on colicin E2 promoter growing in microfluidic growth chambers. Cells express a fluorescent reporter for colicin (*PcolE2\**-mVenus).

***Movie 6.*** Time lapse of MG1655 colicin E2 producing strains with colicin attackers (red) with (top) Low, (middle) WT, and (bottom) High LexA binding strengths on colicin E2 promoter growing in microfluidic growth chambers under the exposure of ciprofloxacin. Cells express a fluorescent reporter for colicin (*PcolE2\**-mVenus).

## Materials and Methods

### Strains, plasmids and genetic constructs

The *E. coli* strains MG1655 and BZB1011 were used as genetic backgrounds in this study. For initial characterization of colicin production dynamics on cell physiology and competitive outcomes, we used BZB1011 harboring a colicin E2 producing plasmid, gifted by Foster group. This strain was transformed with the reporter plasmid pUA66-P*colE2*::gfp KanR^4,30^ and constitutively expressed mTagBFP. To characterize colicinogenic strains carrying modified LexA binding sites, we designed promoter variants based on incorporating the modifications described by Mavridou et al.^30^ We designed two transcription units, first one was the native colicin E2 operon with modified LexA binding sites, the second one was the modified colicin E2 promoter driving the expression of a fast maturing fluorescent protein mVenus^53^, followed by a strong terminator. Both transcription units were assembled using Golden Gate cloning into a p15A origin plasmid conferring kanamycin resistance. The resulting LexA-variant plasmids were transformed into MG1655 strain, which constitutively expressed P*rpsL*-mKate2 for cell tracking. The DNA damage reporter was constructed using Golden Gate assembly. The promoter region of recA (P*recA*) was cloned upstream of a medium-strength ribosome binding site (RBS) driving expression of the fluorescent protein sCFP3A, followed by a strong transcriptional terminator. The P*recA* transcriptional unit was assembled into a low copy pSC101 plasmid backbone carrying a chloramphenicol resistance marker.

### Media and growth conditions

Strain construction was performed in LB medium with appropriate antibiotics when required. Transformants were recovered in SOC medium prior to plating. For planktonic growth, cells were inoculated at 37°C with shaking at 200 rpm in glass tubes with 4 mL medium from a single colony streaked on LB plate, from -80°C stock, within a week of the experiment, unless mentioned otherwise. Strains were stored at -80°C in glycerol stocks. For experiments, cells were grown in EZ Rich Defined medium (EZRDM; TekNova) supplemented with 0.2% glucose. During microfluidic experiments in mother machine devices, fresh medium was provided using peristaltic pumps at speed ∼240 *μ*Lmin^-1^. For experiments in growth chambers, lower flow rates were used to not dislodge cells from the chambers.

### Bulk competition assays

For bulk competition experiments, overnight grown colicin-producing and sensitive strains were mixed at the ratios specified in the Results section and figures, maintaining a final overall starting density equivalent to a 1:50 dilution in 5 mL EZ Rich Defined Medium. Cultures were incubated at 37°C with shaking at 200 rpm to ensure well-mixed conditions. Samples were collected at 2 h and 6 h post-inoculation. Appropriate serial dilutions (10⁻⁵ and 10⁻⁶) were prepared in fresh EZRDM medium and plated on freshly prepared LB agar plates. Plates were incubated overnight at 37°C. Colony counts were obtained by imaging plates using a macroscope in fluorescence channels corresponding to the attacker and sensitive strain, followed by manual quantification of colony counts to determine strain frequencies.

### Microfluidics

Single-cell timelapse imaging was performed using two microfluidic devices: a mother machine device and a spatial competition device with growth chambers. Both devices were fabricated following established protocols^81,82^.

The mother machine chip comprised a main flow channel (25 *μ*m X 80 *μ*m) for providing growth media to cells growing in trenches perpendicular to the main channel. The trenches were 25 *μ*m long with height and width of 1.5*μ*m.

The growth-chamber chip comprised of a similar main flow channel (25 X80 *μ*m) connected to perpendicular growth chambers of dimension 100 *μ*m long, 25 *μ*m wide and 1.4*μ*m high, enabling spatially structured growth of strains. Polydimethylsiloxane (PDMS; 10:1 base-to-curing agent ratio) was poured onto the wafer after degassing and was cured at 95°C for 1.5 h and then peeled off. Inlet and outlet ports were punched prior to bonding. PDMS devices and glass coverslips were plasma-cleaned and bonded, followed by incubation at 95°C for 40 min to strengthen bonding.

For mother machine experiments, stationary phase overnight cultures were loaded into the trenches and continuously supplied with fresh EZRDM. For competitions in chambers, cells were sequentially loaded in lower numbers to not pre-expose sensitive cells to colicin released by colicinogenic cells. The colicinogenic strains were washed thrice with fresh EZRDM media before loading into the microfluidic devices to prevent any carryover of released colicin.

### Microscopy and image analysis

Time-lapse microscopy was performed on a Nikon Ti2 inverted microscope equipped with an Okolab incubation system maintaining the stage at 37°C. Images were acquired every 3 minutes using high-intensity LED or laser illumination, with 100% intensities, and appropriate exposure times optimized for each fluorophore. For the experiments, we used BFP (wavelength = 405 nm), GFP (wavelength = 488 nm), mKate2 (wavelength = 561 nm), Venus (wavelength = 518 nm), and sCFP3A (wavelength = 445 nm). Imaging was performed using a Plan Apo λ 40x Ph2 DM objective. We acquired images with a high pixel density using sCMOS camera (Iris 15). Images were saved as *.nd2* files. Image analysis of mother machine data was performed using BACMMAN^83^ (a Fiji^84^ plugin) followed by custom Python scripts. Preprocessing included drift correction in the x–y plane and alignment of individual growth channels over time. Cell segmentation was performed using the designated segmentation marker channel to delineate cell outlines from background. For analysis of cell fate and lysis dynamics, filamenting cells that exited trenches before the end of the experiment were excluded from analysis. Mean fluorescence intensity was quantified for each segmented cell mask at each time point. Images acquired from growth chamber experiments were analyzed manually to quantify spatial dynamics and strain frequencies over time.

### Analysis

#### Estimating bacterial generation time

To normalize absolute experimental time scales into standardized bacterial generations, single-cell lineage tracking histories were parsed across independent imaging locations and microfluidic channels. The frame indices marking individual cell birth and subsequent division events were extracted. The mean generation time in frames was calculated as

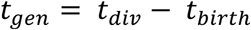

#### Estimating bacterial elongation rate across different colicin promoter expression values

To analyze the relationship between gene expression and cell growth, single-cell tracking data from all acquired positions were pooled. To exclude early transient behaviors and ensure measurements reflected steady-state conditions, data from the first 30 frames were discarded. Raw fluorescence measurements were baseline-corrected by subtracting the global minimum P*colE2-*GFP mean intensity across the dataset from all single-cell fluorescence values. Continuous single-cell lineages were grouped by position, microfluidic channel, and unique lineage identifier. Only established lineages tracked for a minimum of four frames were included in the growth rate analysis. For each individual cell lineage, the specific elongation rate was calculated by fitting a first-degree polynomial (linear regression) to the natural logarithm of the cell length as a function of time (assuming a 3-minute acquisition interval). To account for tracking artifacts or temporary cell shrinkage, any calculated negative growth rates were truncated to zero. The representative gene expression level for each lineage was defined as the median P*colE2*-GFP intensity across its tracked lifespan. To systematically evaluate population-level trends between expression and growth, the continuous single-cell median intensities were discretized into uniform expression bins with a width of 200 arbitrary units (spanning an expression range from 0 to 7000). For each bin, the median fluorescence intensity, median elongation rate, and the standard error of the mean (SEM) of the elongation rate were calculated. Bins lacking sufficient data points were excluded. A weighted linear regression model was fitted to the binned data using non-linear least squares optimization (scipy.optimize.curve_fit), where weights were determined by the SEM of the elongation rate to reduce the influence of highly variable bins. The goodness of fit was quantified using the coefficient of determination (R^2^).

#### Hill coefficient estimation

Single-cell expression traces were categorized into two distinct populations: lysing and non-lysing lineages. *Lysing Lineages*: Lineages that terminated prior to the end of the acquisition period were identified as lysing cells and confirmed manually for lysis events. To determine the P*colE2* expression level responsible for cell death, a 3-frame rolling average was applied to the final 90 frames of the cell’s lifespan, and the maximum smoothed intensity prior to lysis was extracted. *Non-Lysing Lineages*: For cells that survived the observation window, mother cell traces were smoothed using a 3-frame rolling average. To capture natural fluctuations in gene expression that did not result in lysis, local maxima (peaks) in these smoothed traces were detected using a standard peak-finding algorithm (peakutils), employing a relative threshold of 0.3 and a minimum peak distance of 5 frames.

To quantify the relationship between P*colE2*-GFP or P*colE2*-mVenus expression and cell death, we calculated the probability of lysis as a function of fluorescence intensity. Local peaks from non-lysing traces were assigned a binary lysis value of 0, while the terminal maximum intensities of lysing traces were assigned a value of 1. These pooled single-cell events were sorted by intensity and discretized into 20 equally spaced linear bins. For each bin, the mean lysis probability and the standard error of the mean (SEM) were calculated. To model the threshold dynamics of lysis, the binned data were fitted to a Hill equation:

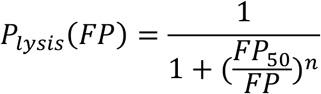

where *P_lysis_*(*FP*)is the probability of lysis at a given fluorescence intensity *FP*, *FP*_50_ represents the threshold expression level required to induce lysis in 50% of the population, and *n* is the apparent Hill coefficient, which dictates the steepness (cooperativity) of the lysis response. Non-linear least-squares optimization was used to estimate the *FP*_50_ and *n* parameters. Data visualization, including probability density functions (PDF) and fitted dose-response curves, was generated using Matplotlib library with custom python scripts.

#### Quantification of intensity and lysis events in spatial competition

Initial image processing was performed using Fiji. To isolate the imaging channels for segmentation marker and P*recA* expression in sCFP3A channel, the original multi-channel Z-stack/time-series was decomposed using the Split Channels function.

To facilitate localized intensity analysis, a custom ImageJ Macro (IJM) was utilized to partition the image into a grid of size 12 pixel by 12 pixel. The macro calculated the width (w) and height (h) of the active image and programmatically generated rectangular regions of interest, which were then added to the ROI Manager.

*Plugin>New>Macro Macro Code:*

*getDimensions(w, h, channels, slices, frames);*

*cellSize = 12;*

*gridX = floor(w / cellSize);*

*gridY = floor(h / cellSize);*

*roiManager(“reset”);*

*for (y = 0; y < gridY; y++) {*

*for (x = 0; x < gridX; x++) {*

*makeRectangle(x * cellSize, y * cellSize, cellSize, cellSize);*

*roiManager(“add”);*

*}*

*}*

*roiManager(“Show All”);*

Keep the Tif files open for which you run the code.

To extract the intensity, we used Analyze > Set Measurements, with Mean Gray Value selected as the primary metric. The ROIs generated in the previous step were applied to the entire temporal stack using the Multi-Measure tool within the ROI Manager. To maintain data structure, the “Measure all slices” and “One row per slice” options were enabled. This resulted in a formatted results table where each row corresponds to a specific time point (t) and each column corresponds to a specific grid coordinate. The resulting intensity-over-time data was exported as a .csv file. Final plotting was performed using custom Python code.

#### Competitive advantage computation

Ratio of attacker to sensitive strain relative to that at time =0.

#### Analysis of lysis events in chamber

The lysis events were mapped by manually inspecting the tiff files.

#### Growth, Elongation, and Dilution-Corrected Promoter Activity

Interdivision times were extracted from continuous single-cell lineages during steady-state pre-treatment conditions to calculate the mean generation time (Td). The specific growth rate (μ) for the cellular population was subsequently defined as μ= ln(2)/Td. To capture dynamic morphological changes, the instantaneous single-cell elongation rate was computed as the temporal derivative of the natural logarithm of cell length, d(lnL)/dt. To quantify transcriptional dynamics independent of passive cell growth and fluorophore dilution, Promoter Activity (PA) was calculated from the smoothed fluorescence intensity (I). We computed PA by taking the numerical temporal derivative of the fluorescence signal and adding a growth-dilution correction term, defined as PA= dI/dt+μI.

#### Signal Processing and Transcriptional Peak Classification

To distinguish stochastic transcriptional fluctuations from deterministic commitment events, raw fluorescence intensities for the fluorescent reporters were subjected to signal processing. Single-cell fluorescence trajectories were first smoothed using a centered 5-frame moving average to minimize frame-to-frame imaging noise. To account for basal autofluorescence and global background drift, each trace was locally baseline-corrected by subtracting the 1st percentile intensity specific to that individual cell’s trajectory.

Algorithmic peak detection (via the SciPy find_peaks function in Python) was then applied to the smoothed, baseline-corrected traces. Transcriptional peaks were identified requiring a minimum temporal separation of 5 frames and a defined prominence threshold tailored to the specific reporter fluorophore to ensure only genuine transcriptional bursts were captured.

### Model for competition in a well mixed environment

Initial condition for attackers (A) and susceptibles (S) mixed in ratio *r*: 1 − *r* is given as:

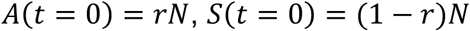

Growth rate for A and S is given by *μ_A_* = *μ_S_* = 1.63 ℎ^−1^. *θ* is the accumulated toxin level in lysis case that leads to cell growth stalling and subsequent lysis, incurring the maximal burden at this concentration.

Susceptible population

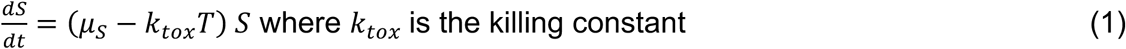

Attacker population

Lysis mode:

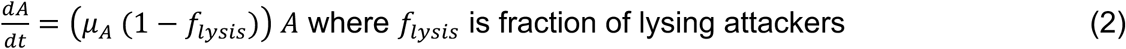

Secretion mode:

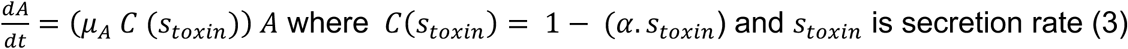

Toxin production

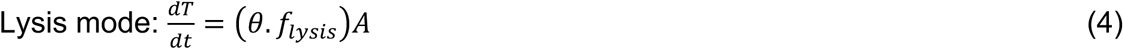

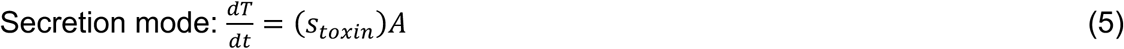

We simulate for different values of *s_toxin_* in secretion mode and associated values of *f_lysis_* in lysis mode when growth penalty is equalized by equating equations 2 and 3,

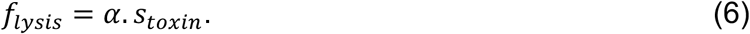

When 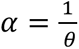, growth penalty and toxin flux are equalized and hence there is no difference in attacker advantage between lysis and secretion modes.

The ratio in toxin production when growth penalty is equated gives:

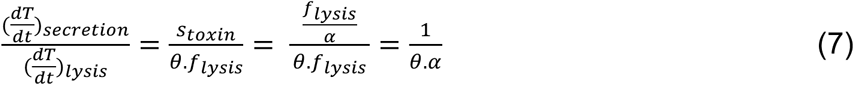

When 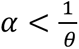 toxin produced by secretion case is higher than toxin produced by lysis case. And toxin production is higher in lysis case when 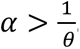.

Simulation outcomes are given by the following:

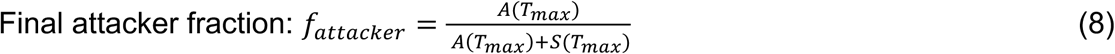

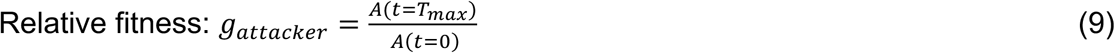

Simulation Conditions: Simulations are initialized at varying starting attacker ratios (*r*), ranging from 0.001 to 0.1. The total initial simulated population was fixed at *N*=10,000 cells, and the initial toxin concentration *T*_0_ = 0. The system of equations was solved numerically using a forward Euler integration scheme with a discrete time step *dt* = 0.01 over a total simulation time *T_max_* = 6 ℎ.

To rigorously assess the model’s sensitivity to estimate lysis threshold *θ*, we systematically scan parameter for *θ*. Twenty-five logarithmically spaced *θ* values spanning four orders of magnitude are evaluated. For each discrete value of *θ*, the bounded optimization of toxin production is repeated for experimental values of well mixed competition for all three variants. The aggregate MSE (mean squared error) across all variants is calculated for *θ* to generate a global goodness-of-fit profile and empirically identify the optimal *θ* that best captures the population dynamics of the experimental data. The lowest error was for *θ* ≥1000

To assess whether *θ* yields agreement between model predictions and experimental values, we compare *f_attacker_* from simulations with experimental *f_attacker_* to predict the lysis fraction *f_lysis_*, for the distinct LexA binding variants (LexA-low, WT, and LexA-high). Fitting is performed by minimizing the MSE between the experimentally observed *f_attacker_* and the simulated *f_attacker_*. The optimization is executed using the minimize_scalar function from the scipy.optimize library, employing a bounded search algorithm constrained within the interval from 10^-6^ to 500. Upon convergence, the variant-specific lysis fraction is computed as shown in Figure 3B.

### Model for spatial competition using lattice model

The simulation is performed in a 2D regular square lattice of size H X W (10 X 40 cell lengths). Space is discretized into grid cells, each capable of holding a maximum of one bacterial cell. The simulation proceeds in discrete time steps, where each step represents a unit of biological time equivalent to a minute.

*Agent dynamics:*

Bacteria are modeled as discrete agents belonging to one of two competitive strains: an “Attacker” strain (employing either a lysis or secretion strategy) and a “Target” (susceptible) strain.

At time t, a cell exists in the following states and divides via a timer with division interval *t_div_* (= 24 min).

0: Empty; 1: Attacker (non-producer); 2: Susceptible; 3: Attacker (sacrificial producer)

Upon division, a daughter cell is placed in a neighboring grid cell determined by the mother cell’s orientation vector. If the intended target grid cell for the daughter is unoccupied, the daughter cell is placed there. If the target cell is occupied, the model simulates localized mechanical shoving to accommodate the new cell. The entire column of cells along the y-axis (from the target coordinate up to the upper boundary) is shifted forward by one grid unit to create empty space, pushing the outermost cells outward.

In the secretion case, attacker cells show inhibition of growth rate as described in Eq 3.

*Toxin production:*

Lysis mode:

In the lysis strategy, individual attacker cells undergo stochastic, catastrophic lysis to release a large bolus of toxin. At each time step, an attacker cell has a fixed probability *lysis* of triggering the lysis program.

Once triggered, the cell ceases to divide, accumulates toxin for a brief period, and then bursts, releasing a fixed quantity of toxin, *θ*, into its local grid environment. The cell agent is then removed from the grid.

Secretion mode: Each attacker secretes *s_toxin_* at each time point.

*Toxin diffusion and decay:*

The spatio-temporal dynamics of the extracellular toxin concentration field evolve as follows:

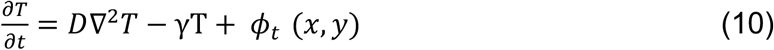

where D is toxin diffusivity and *φ_t_* is the toxin source (lysis/secretion) and γ is first-order toxin decay term due to flow in the chambers.

To ensure numerical stability when solving the diffusion PDE on the discrete lattice, the PDE is numerically integrated using a forward Euler finite difference scheme with *n* =20 temporal substeps per cellular time step. Spatial derivatives are computed using a 2D discrete Laplacian convolution kernel with reflective boundary conditions (∇*C*. *n* = 0 at the grid boundaries).

The discretized update rule for substepping is given by:

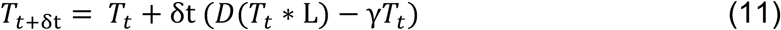

where δt = 1/*n* and L is the 3X3 Laplacian kernel.

Toxin source:

Secretion Mode: Attacker releases toxin continuously

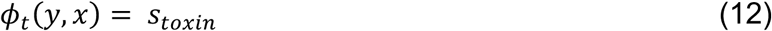

Lysis Mode: Sacrificial attacker produces a single burst of accumulated toxin *θ*

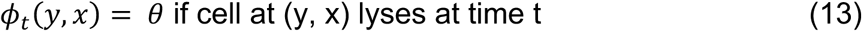

Sensitive cell death: Sensitive cells (cell state=2) are removed if the local toxin concentration exceeds a lethal threshold *T_let_*_ℎ*al*_.

For simulating cell movement in the grid, we assume each cell has an orientation *ζ*. At each time step, *ζ* undergoes rotational diffusion:

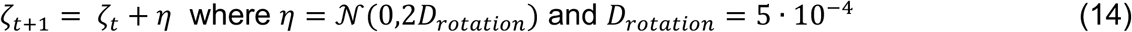

Again, to equate growth penalty in lysis and secretion mode: *f_lysis_* = *α*. *s_toxin_* as in Eq 6. We simulate different values of *s_toxin_* in secretion mode and associated values of *f_lysis_* in lysis mode where 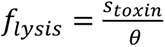 where *f_lysis_*<0.5.

Simulation outcomes:

Let N be the total number of lattice sites. Let *c_s_*(*y*, *x*) be the site occupied by state S in the cell at (y,x). *L* = ∑_(*y*,*x*)_ *c_s_*_∈(1,2,K)_(*y*, *x*) is the total number of lattice sites occupied by cells.

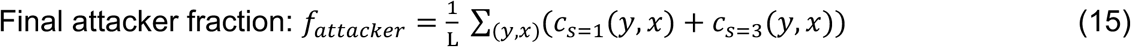

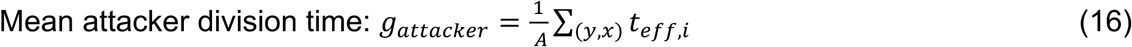

where A is the number of attackers and *t_eff_*_,*i*_ is the toxin-inhibited interval for cell i. The concentration T at distance r and time t after *θ* in two dimension is given as:

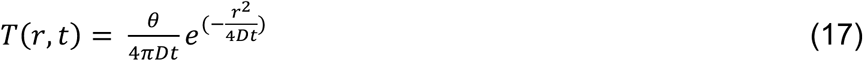

The killing zone expands as toxin diffuses and then fades as the concentration drops.

For any distance r, the maximum concentration occurs at time 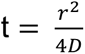.

Substituting this in the above equation we get

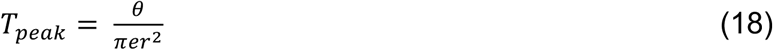

Now setting *T_peak_*(*R_max_*) = *T_let_*_ℎ*al*_ and solving for *R_max_*, we get:

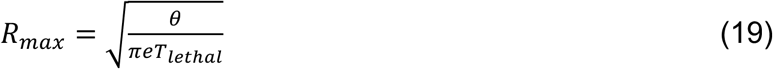

So, *R_max_* α √*θ*

We substitute killing radius of about 5 cells, which gives *R_max_*∼5 at which a cell could die in the simulations which yields the following relationship.

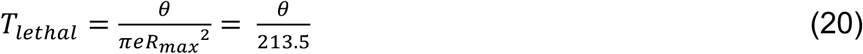

Susceptible (target) cells monitor the local toxin concentration in their resident grid cell. If the environmental toxin concentration meets or exceeds a critical lethal threshold, *T_let_*_ℎ*al*_, the susceptible cell is immediately killed and removed from the grid. Simulations are initialized with a localized band of cells at the bottom boundary (the first 5 rows) to simulate a growing front. The initial population is randomly mixed with a predefined initial ratio *r* of attacker cells to target cells and is kept the same for comparing the secretion and lysis strategy.

---

**Figure S1:**
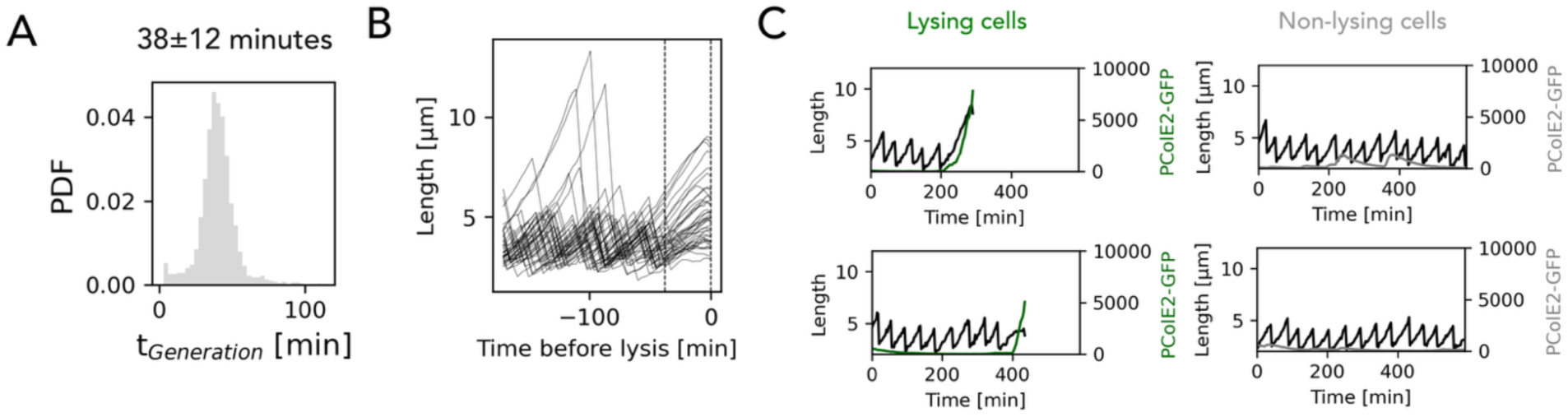
(A) Distribution of generation time of colicin producing strain in Figure 1. (B) Traces of length for lysing attackers followed from 150 minutes before lysis to assess length changes that preceded lysis, dashed line represents average division time. (C) Length (black) and *PcolE2-GFP* (green for lysing or grey for alive) expression traces of two representative lysing (green, left) and non-lysing (grey, right) mother cells.

**Figure S2:**
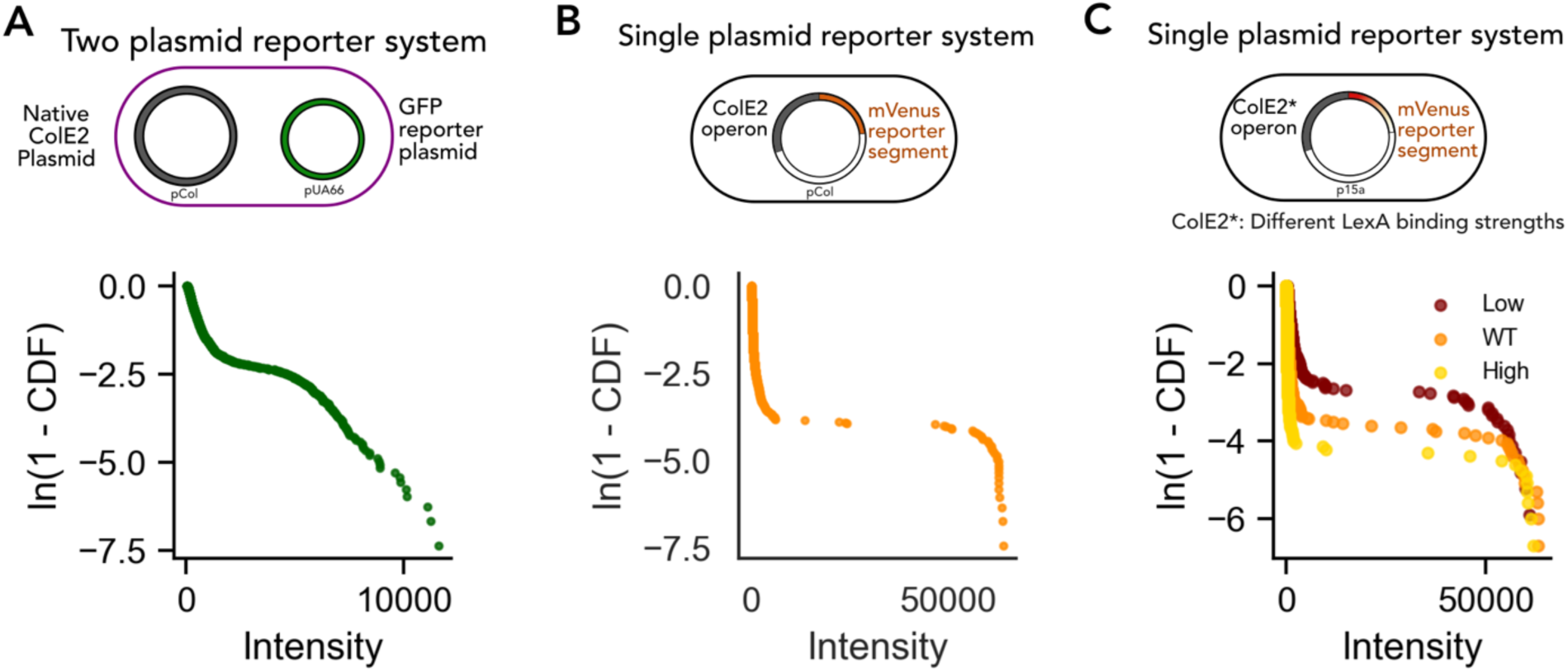
Bimodal distribution of peak intensities of colicin expression. (A) (top) Schematic of the two-plasmid reporter system with native colicin E2 plasmid and PcolE2-GFP reporter system. The native pColE2 plasmid encodes the ColE2 toxin, immunity protein (ImmE2), and lysis machinery, while a second plasmid carries a PcolE2-GFP reporter to monitor toxin production dynamics. (bottom) Distribution of peak intensity of colicin expression (P*colE2*-GFP) shown as complementary cumulative distribution plotted as ln(1-CDF). (B) (top) Schematic of the single-plasmid reporter system, in which an mVenus reporter is placed under the control of the native Pc*olE2* promoter, enabling simultaneous monitoring of colicin expression and cell fate from the native colicin plasmid. (bottom) Distribution of peak intensity of colicin expression (P*colE2*-mVenus) shown as complementary cumulative distribution plotted as ln(1-CDF). (C) (top) Schematic of the single-plasmid reporter system, in which an mVenus reporter is placed under the control of the Pc*olE2\** promoter with modified LexA-binding-site, alongside transcription unit of colicin under the control of Pc*olE2\** promoter with modified LexA-binding-site. (bottom) Distribution of peak intensity of colicin expression (P*colE2\**-mVenus) shown as complementary cumulative distribution plotted as ln(1-CDF). (LexA-binding-strength: low (maroon), WT (orange) and high (yellow))

**Figure S3:**
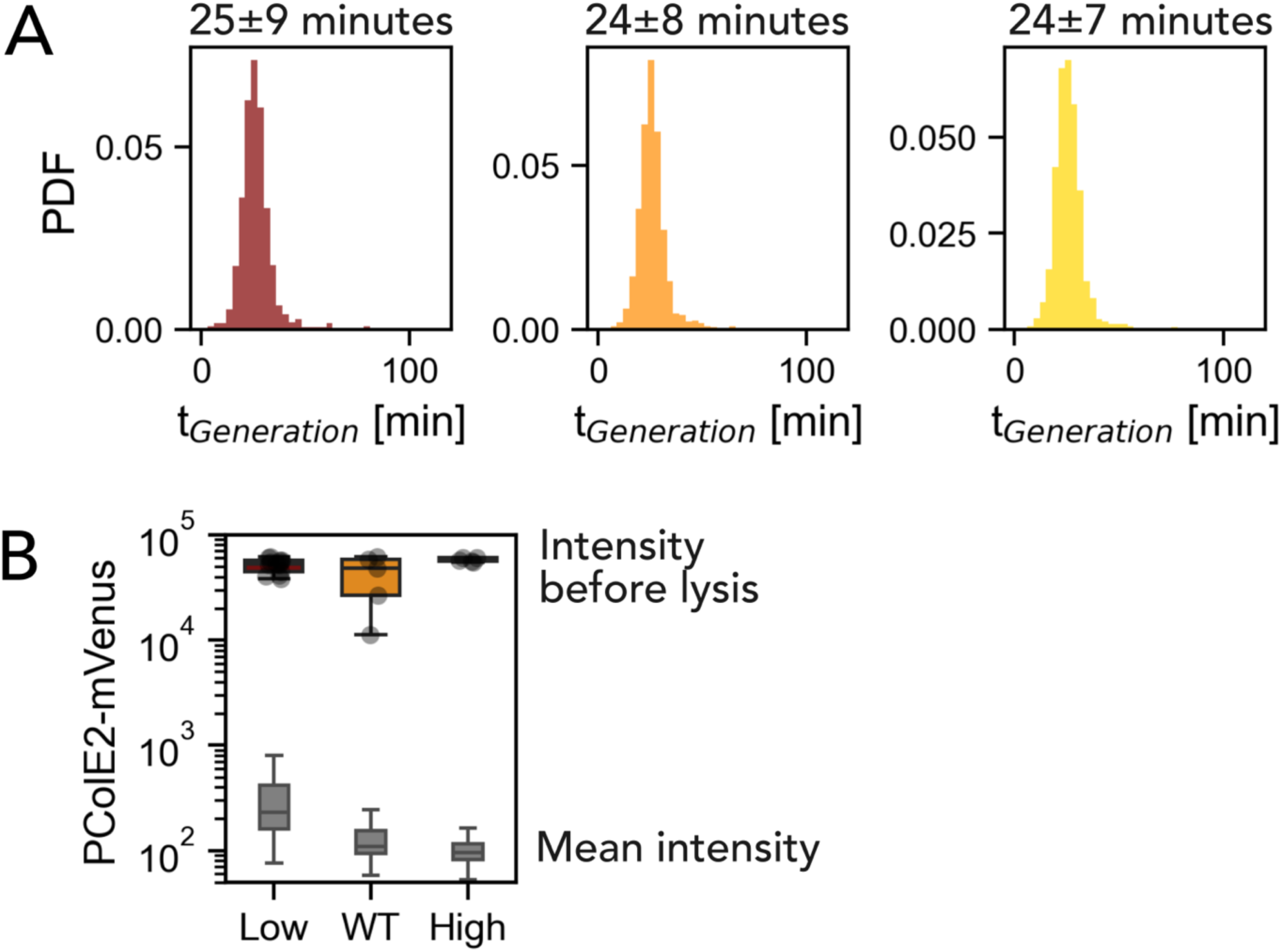
(A) Distribution of generation time for (left, maroon) low, (center, orange) WT and (right, yellow) high LexA binding strains in Figure 2. (B) Box plot representing average *PcolE2\**-mVenus intensity before cell lysis (colored) and intensity of all cells (grey) for low (maroon), WT (orange) and high (yellow) LexA binding strains (error bar = standard deviation).

**Figure S4:**
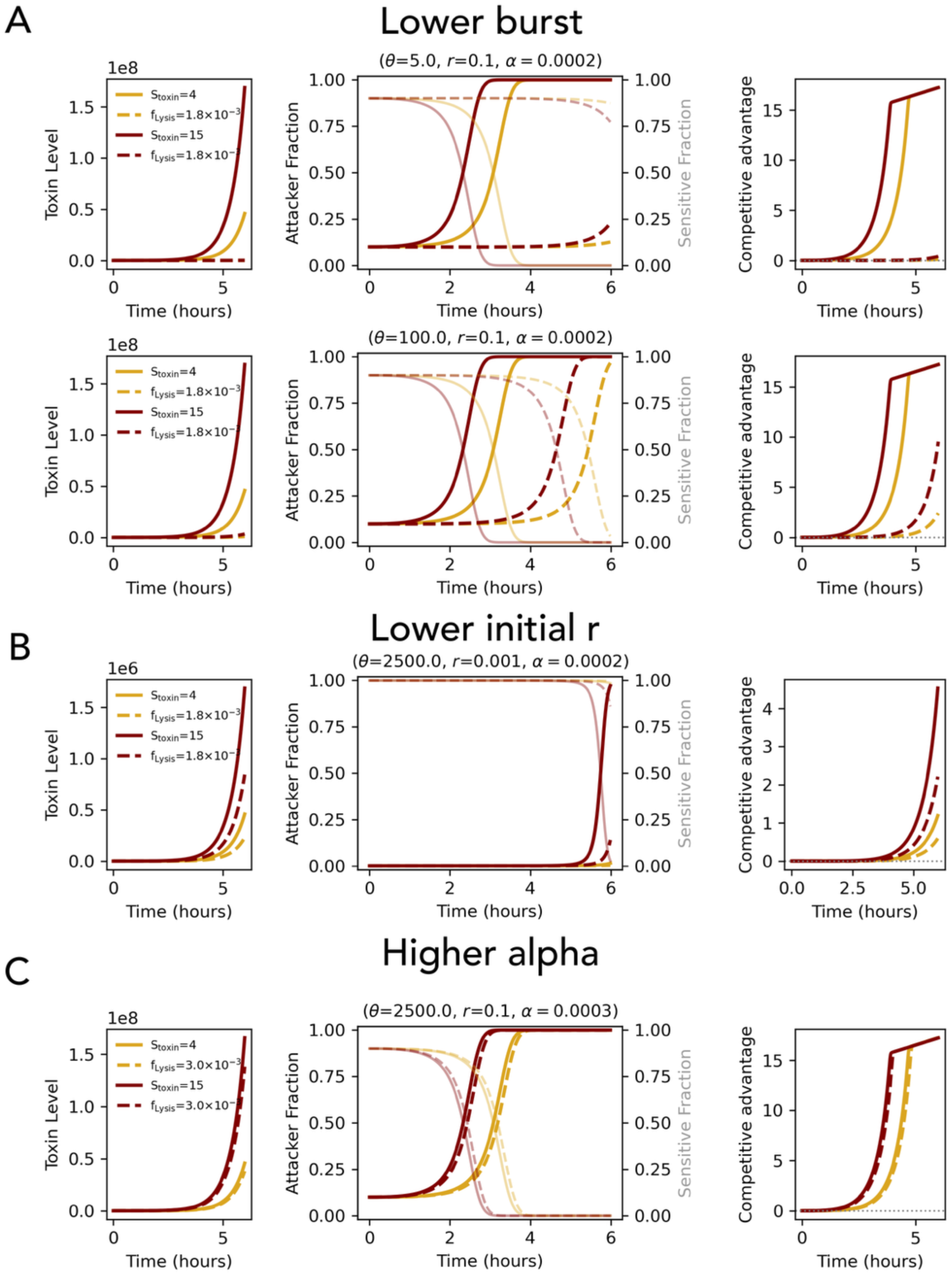
(A) Traces comparing secretion (bold, uniform labor) versus lysis (dashed, division of labor) mode of competition between sensitive and attacker strains in well mixed environments. Results for low (yellow) and high toxin production (maroon) with different lysis threshold values (top: *θ* = 5 and bottom *θ* = 100). Competitions start with initial ratio *r*=0.1. (left) Toxin level accumulated over time. (center) Attacker (dark) and sensitive (light) fraction over time during competition. (right) Competitive advantage over time. (B) Competitions similar to those in panel A but starting with initial ratio *r*=0.001. (left) Toxin level accumulated over time. (center) Attacker (dark) and sensitive (light) fraction over time during competition. (right) Competitive advantage over time. (C) Competitions similar to panel A and B but with higher growth penalty *α*. Competitions start with initial ratio *r*=0.1. (left) Toxin level accumulated over time. (center) Attacker (dark) and sensitive (light) fraction over time during competition. (right) Competitive advantage over time.

**Figure S5:**
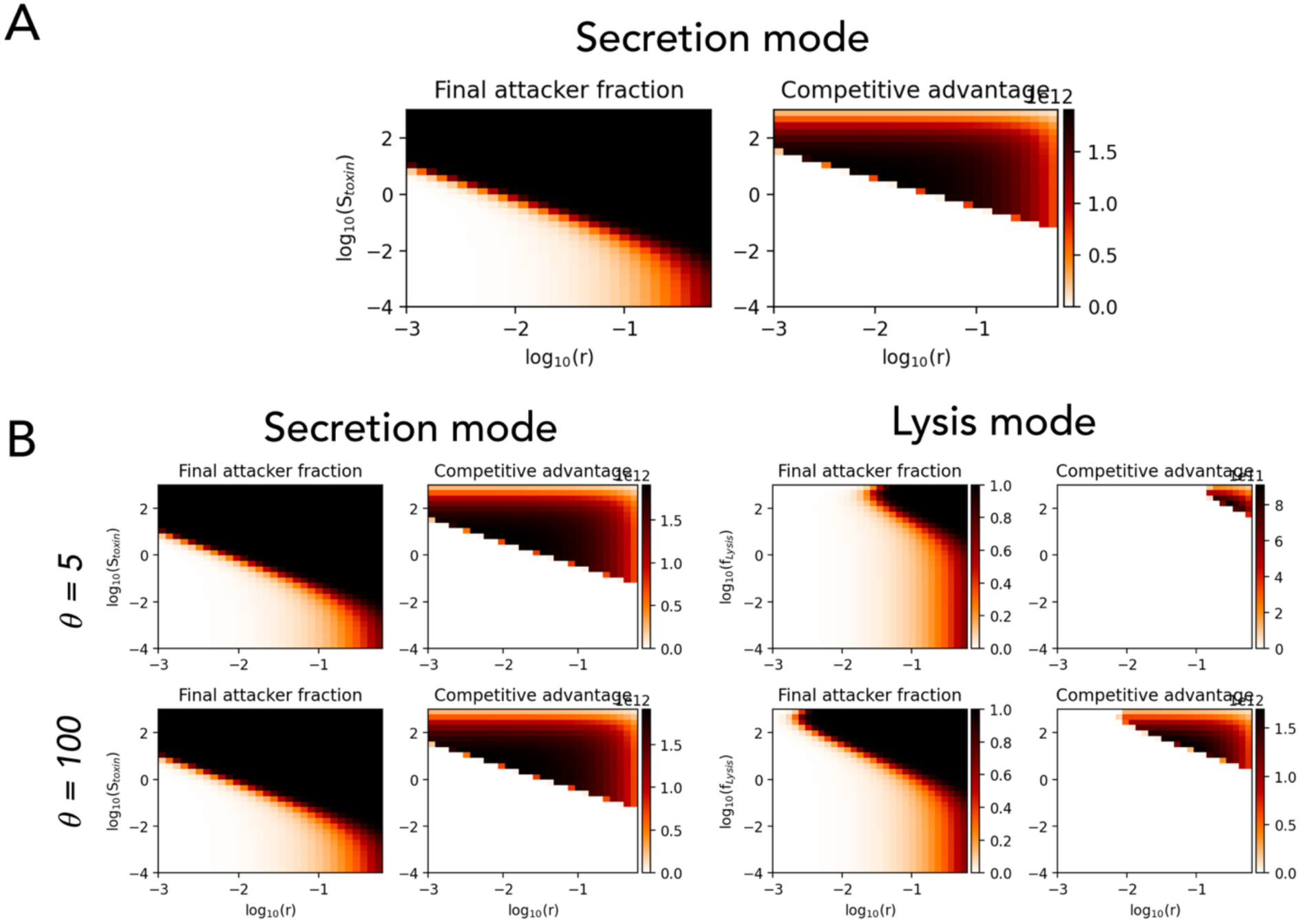
(A) Heatmap of (left) final attacker fraction and (right) competitive advantage for well mixed competition in secretion mode with S_toxin_ values equated against those of f_lysis_ in Figure 3C, simulations ran based on Figure 3A. (B) Heatmap of final attacker fraction and (right) competitive advantage for well mixed competition in (left) secretion and (right) lysis mode with equated growth penalty for different toxin accumulated before lysis (*θ*) (top: *θ* = 5 and bottom: *θ* = 100).

**Figure S6.**
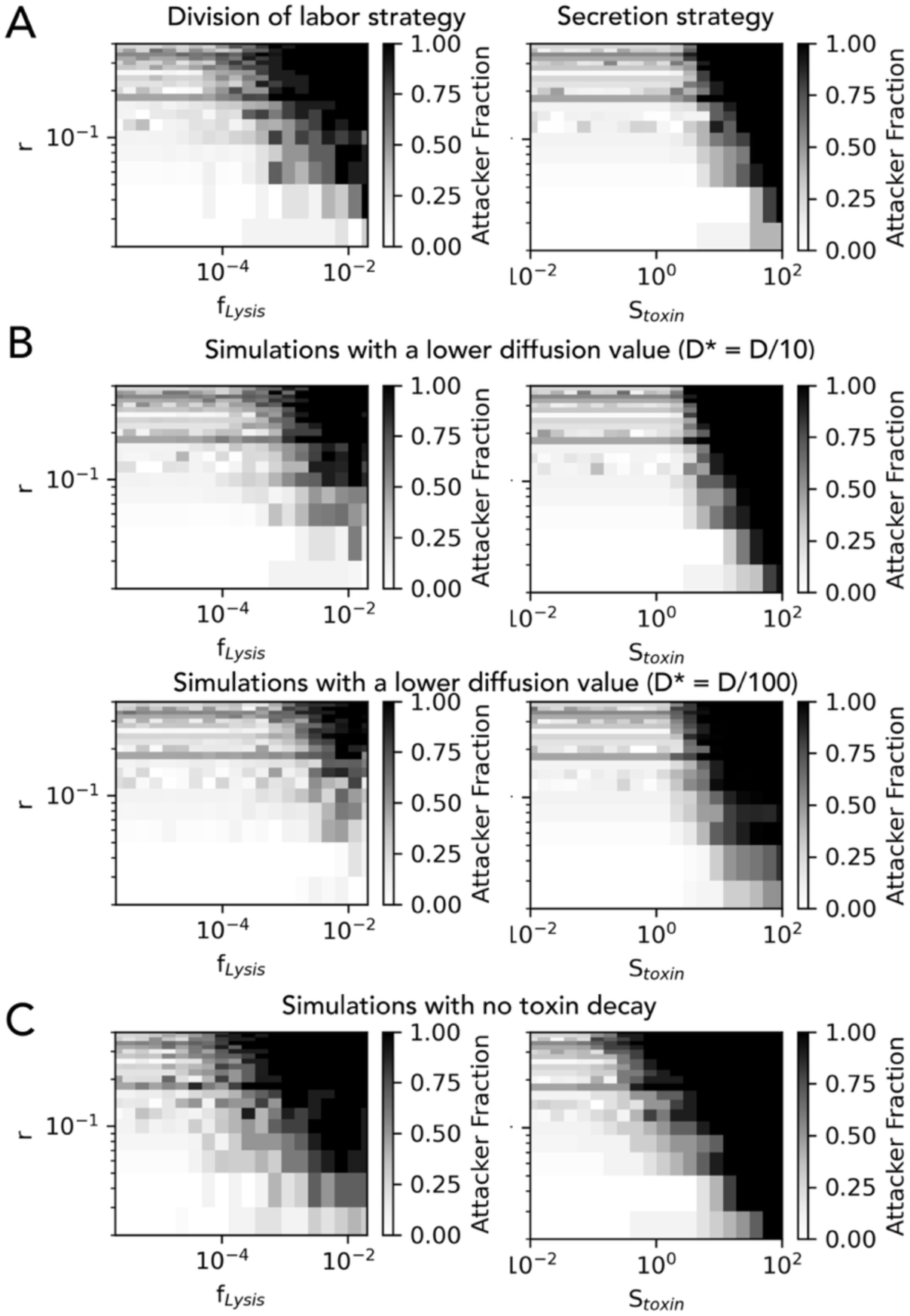
(A) Heatmap of final attacker fraction for (left) lysis strategy (division of labor) and (right) secretion strategy (uniform labor) for spatial competition with high diffusion values and high toxin decay. Darker colors showing areas where attackers dominate. (B) Heatmap of final attacker fraction for (left) lysis (division of labor) strategy and (right) secretion strategy (uniform labor) for spatial competition with lower diffusion values and high toxin decay. Darker colors showing areas where attackers dominate. (C) Heatmap of final attacker fraction for (left) lysis (division of labor) strategy and (right) secretion strategy (uniform labor) for spatial competition with high diffusion values and no toxin decay. Darker colors showing areas where attackers dominate.

**Figure S7.**
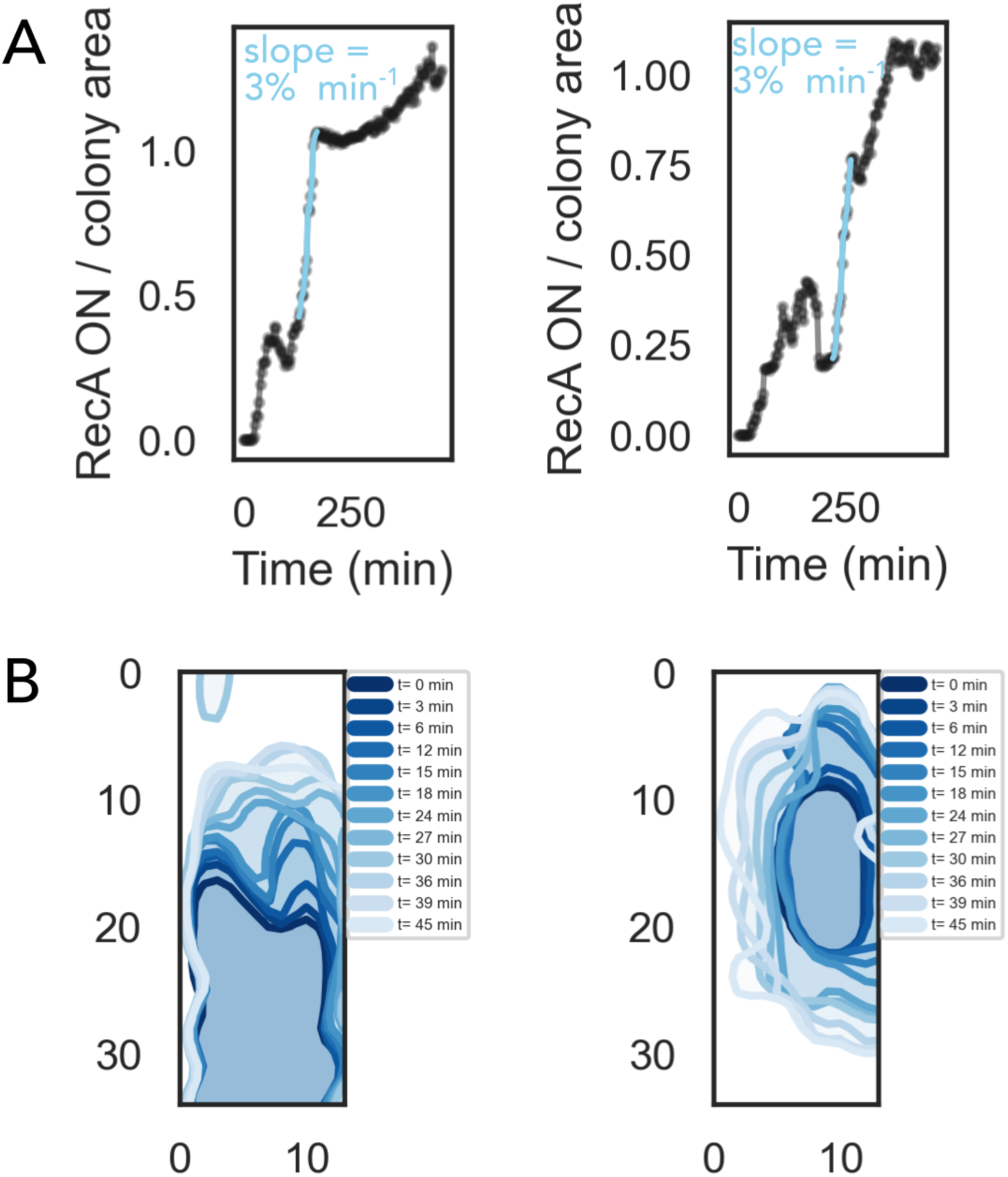
(A) Fraction of colony that is occupied by sensitive cells and exhibits P*recA* ‘ON’ over time for two chambers competing a colicin producing and sensitive strain, the blue points show the time when the DNA-damage by colicin released by lysing attacker damages the sensitive cells. (B) (The spatio-temporal spread of P*recA* ‘ON’ in chamber area occupied by sensitive cells during the time depicted in blue in panel A for two chambers.

**Figure S8.**
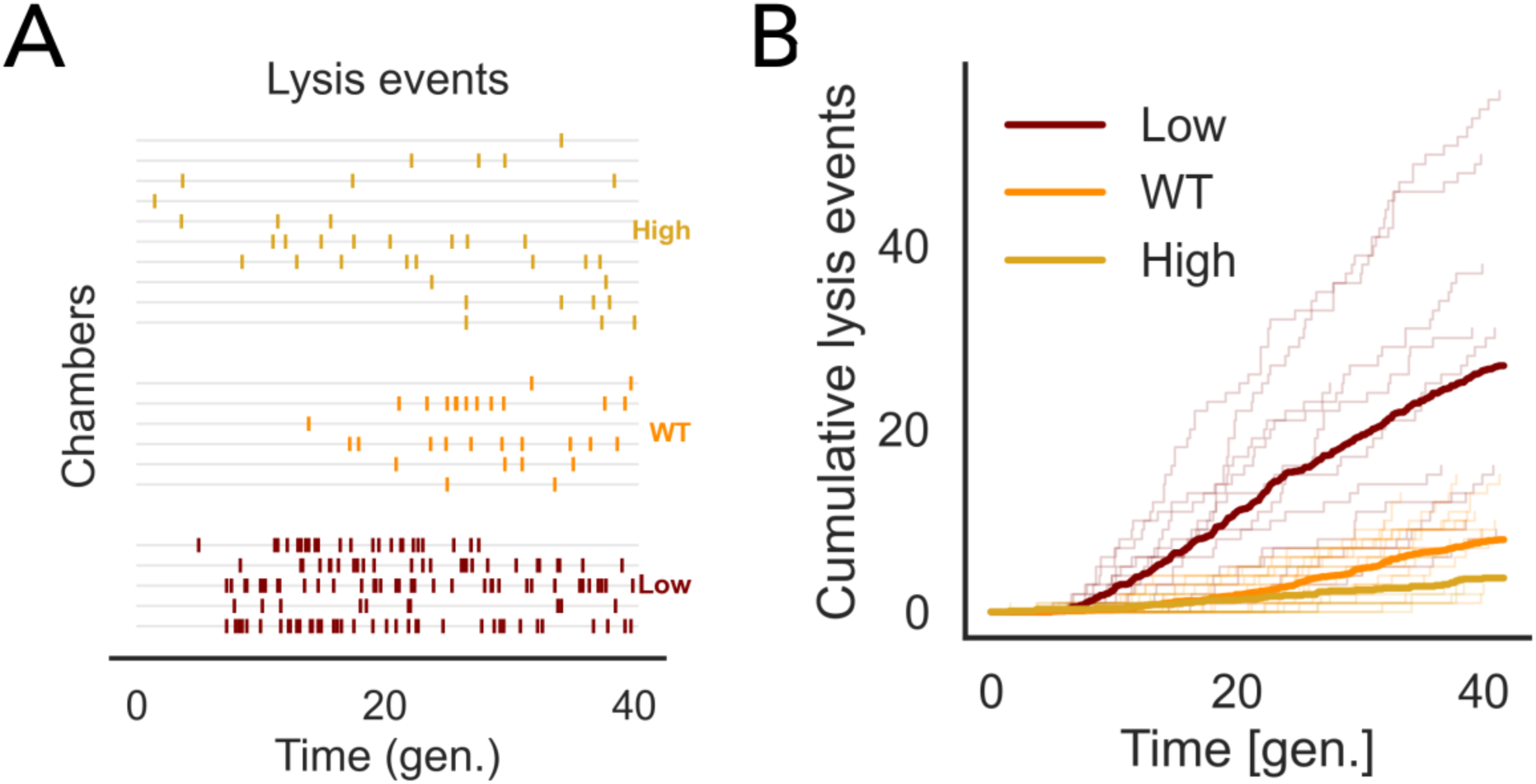
(A) Traces depicting lysis events in individual chambers for (top) high, (center) WT, and (bottom) Low LexA binding strains. (B) Cumulative lysis events over time in each chamber (thin lines, mean depicted by thick lines) for (yellow) high, (orange) WT, and (maroon) Low LexA binding strain.

**Figure S9.**
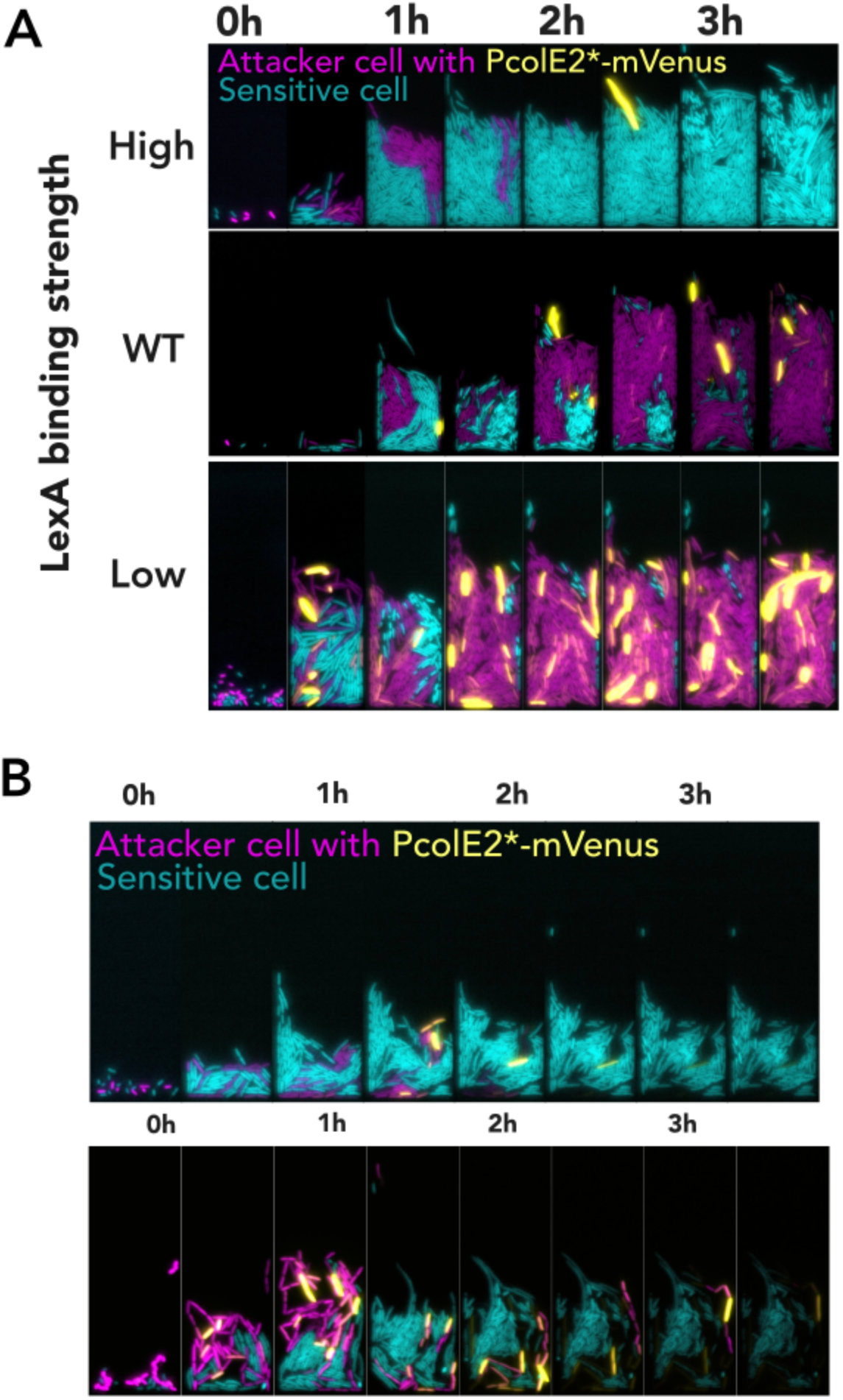
High colicin production risks lineage collapse during spatial warfare (A) Kymographs of competitions in microfluidic chambers using the attackers (magenta) with High, WT, and Low LexA binding strengths against sensitive cells (cyan). (B) Representative time-lapse images showing population collapse in a chamber initiated with the high-colicin producing (Low LexA binding site) strain (magenta) competing with sensitive strain (cyan).

## Notes

### Competing Interest Statement

The authors have declared no competing interest.

